# Phase synchrony between prefrontal noradrenergic and cholinergic signals indexes inhibitory control

**DOI:** 10.1101/2024.05.17.594562

**Authors:** Yuxiang (Andy) Liu, Yuhan Nong, Jiesi Feng, Guochuan Li, Paul Sajda, Yulong Li, Qi Wang

## Abstract

Inhibitory control is a critical executive function that allows animals to suppress their impulsive behavior in order to achieve certain goals or avoid punishment. We investigated norepinephrine (NE) and acetylcholine (ACh) dynamics and population neuronal activity in the prefrontal cortex (PFC) during inhibitory control. Using fluorescent sensors to measure extracellular levels of NE and ACh, we simultaneously recorded prefrontal NE and ACh dynamics in mice performing inhibitory control tasks. The prefrontal NE and ACh signals exhibited strong coherence at 0.4-0.8 Hz. Although inhibition of locus coeruleus (LC) neurons projecting to the PFC impaired inhibitory control, inhibiting LC neurons projecting to the basal forebrain (BF) caused a more profound impairment, despite an approximately 30% overlap between LC neurons projecting to the PFC and BF, as revealed by our tracing studies. The inhibition of LC neurons projecting to the BF did not diminish the difference in prefrontal NE/ACh signals between successful and failed trials; instead, it abolished the difference in NE-ACh phase synchrony between successful and failed trials, indicating that NE-ACh phase synchrony is a task-relevant neuromodulatory feature. Chemogenetic inhibition of cholinergic neurons that project to the LC region did not impair inhibitory control, nor did it abolish the difference in NE-ACh phase synchrony between successful or failed trials, further confirming the relevance of NE-ACh phase synchrony to inhibitory control. To understand the possible effect of NE-ACh synchrony on prefrontal population activity, we employed Neuropixels to record from the PFC during inhibitory control. The inhibition of LC neurons projecting to the BF not only reduced the number of prefrontal neurons encoding inhibitory control, but also disrupted population firing patterns representing inhibitory control, as revealed by a demixed principal component (dPCA) analysis. Taken together, these findings suggest that the LC modulates inhibitory control through its collective effect with cholinergic systems on population activity in the prefrontal cortex. Our results further indicate that NE-ACh phase synchrony is a critical neuromodulatory feature with important implications for cognitive control.

## Introduction

Whether “biting one’s tongue” at the Thanksgiving table during a political conversation or laying off a pitch out of the strike zone, the ability to inhibit inappropriate behavior to achieve a specific goal is a critical element of our executive function. In general, inhibitory control enables animals to suppress their impulsive behavior until conditions are appropriate, preventing undesired or sub-optimal outcomes^1^. Impulsivity is a complex neuropsychiatric trait and is often referred to as the tendency of rapid but often premature actions without foresight. Impulsive behavior is widely believed to result from impaired “top-down” inhibitory control. It is a hallmark of several major clinical conditions, including substance abuse disorder, attention deficit hyperactivity disorder (ADHD), and antisocial personality disorder^2^. A growing body of research suggests that the prefrontal cortex is a central node in the brain’s impulsivity network, playing a crucial role in inhibitory control ^3–5^. Moreover, several neurotransmitter systems profoundly influence cognitive functions, including inhibitory control ^6–12^. The dopaminergic system has long been implicated in impulse control. This is due to the dramatic therapeutic efficacy of amphetamine, a dopamine agonist, and methylphenidate, a dopamine and norepinephrine reuptake inhibitor, in treating impulsivity symptoms in ADHD patients. More recently, several lines of evidence from preclinical and clinical studies have indicated the involvement of the noradrenergic and cholinergic systems in inhibitory control^7–12^. For example, Robinson *et al.* ^8^ found that administering Atomoxetine, a selective norepinephrine reuptake inhibitor, significantly improved the impulse control of rats across various behavioral tasks measuring impulsivity.

Norepinephrine (NE), along with acetylcholine (ACh), are two essential neurotransmitters in the brain. NE/ACh, released from the axon terminals of noradrenergic/cholinergic neurons, exerts an effect on noradrenergic and cholinergic receptors mainly through volume transmission to influence a variety of sensorimotor and cognitive functions^13–15^. The brainstem noradrenergic nucleus, the locus coeruleus (LC), provides the primary source of norepinephrine input to the entire forebrain^16,17^. The LC modulates various brain functions through its diffuse projections throughout the brain^18–25^. Similarly, cholinergic neurons within the basal forebrain region are the primary source of cholinergic input to the cortex^26^. The prefrontal cortex (PFC) is heavily innervated by cholinergic and noradrenergic systems. PFC neurons co-express adrenergic and cholinergic receptors, suggesting that the two neurotransmitters may engage competing intracellular signaling pathways ^27–29^. However, little is known about the dynamics of NE and ACh in the prefrontal cortex during inhibitory control. Moreover, although previous work utilizing anatomic tracing, pharmacological manipulation and modeling has suggested the role of interaction between the NE and ACh systems in modulating cognitive functions ^30,31^, the extent to which the NE-ACh interaction modulates prefrontal population activity and, in turn, inhibitory control remains poorly understood.

To address these questions, in the present study, we simultaneously measured extracellular NE and ACh levels using florescent GRAB_NE_ and GRAB_ACh_ sensors in mice performing an inhibitory control task, which required the mice to suppress impulsive licking, to uncover the dynamics of prefrontal NE and ACh during inhibitory control. Here, we show that the phase relationship between prefrontal NE and ACh signals was dynamic during inhibitory control, with the two signals more likely being in-phase. Chemogenetic inhibition of LC neurons that project to the basal forebrain region reduced behavioral performance to a chance level. Surprisingly, this manipulation abolished the difference in NE-ACh phase synchrony, but not the difference in the NE/ACh signals between successful and failed trials. Chemogenetic inhibition of cholinergic neurons projecting to the LC did not alter prefrontal NE-ACh phase synchrony, nor did it affect inhibitory control performance. Subsequent Neuropixels recordings from the prefrontal cortex confirmed that inhibition of LC neurons that project to the basal forebrain region also disrupted population dynamics representing inhibitory control, suggesting a modulatory effect of NE-ACh phase synchrony on neural activity. Retrograde tracing revealed distinct subgroups of LC neurons projecting to the PFC or the basal forebrain, with approximately 30% overlap between the two groups. Taken together, these results indicate that prefrontal NE-ACh phase synchrony is a novel neuromodulatory feature that indexes neuromodulation of population activity mediating inhibitory control.

## Results

### Correlated fluctuations of NE and ACh levels in the prefrontal cortex

To investigate the interaction between the noradrenergic and cholinergic signals in inhibitory control, we used AAV vectors to express genetically encoded NE and ACh fluorescent biosensors GRAB_NE_ and GRAB_ACh_ in the prefrontal cortex of head-fixed mice (**Figure 1a, b**; one biosensor in each hemisphere, randomly assigned, GRAB_NE_ in the left prefrontal cortex of 9 mice out of 19 mice). During the initial shaping period, we observed a transient increase of both NE and ACh levels following the random delivery of sweet water, confirming the ability of these biosensors to index behavior (**Figure 1c, d**). Furthermore, the amplitude of the transient ACh increase elicited by sweet water rewards (and possibly consequent licking activities) was comparable to that of transient NE responses (**Figure 1e**, p=0.22; Wilcoxon signed-rank test). We also found a significant difference in the peak response latency between NE and ACh responses. (**Figure 1e**, 1p<0.03; Wilcoxon signed-rank test). The level of both NE and ACh in the brain fluctuated spontaneously (**Figure 1f**). However, the fluctuation of the neurotransmitters was correlated. Cross-correlation analysis revealed a positive correlation between NE and ACh with a peak correlation coefficient of 0.331±0.041, significantly greater than 0 (p<1.3e-4, Wilcoxon signed-rank test). Consistent with previous work, NE dynamics preceded ACh dynamics by 0.03±0.01 seconds (p<0.0056, Wilcoxon signed-rank test, **Figure 1g**) ^32^. Coherence analysis revealed that the two neuromodulatory signals exhibited a maximum correlation at a frequency range of 0.4-0.8 Hz, indicating a strong interplay at this frequency band (**Figure 1h**). Since we simultaneously recorded GRAB_NE_ and GRAB_ACh_ signals, one in the PFC of each hemisphere, we first examined whether the GRAB signal in one hemisphere is a good estimate of the same GRAB signal in the other hemisphere. We simultaneously recorded GRAB_NE_ or GRAB_ACh_ signals from the PFC of both hemispheres and computed cross-correlograms using band-pass filtered (0.4– 0.8 Hz) GRAB signals. For both GRAB signals, the correlation coefficient at lag 0 was approximately 0.85 (**Supplemental figure 1a**), confirming that the GRAB signal in one hemisphere serves as a reliable estimate of the same GRAB signal in the other hemisphere.

**Figure 1.**
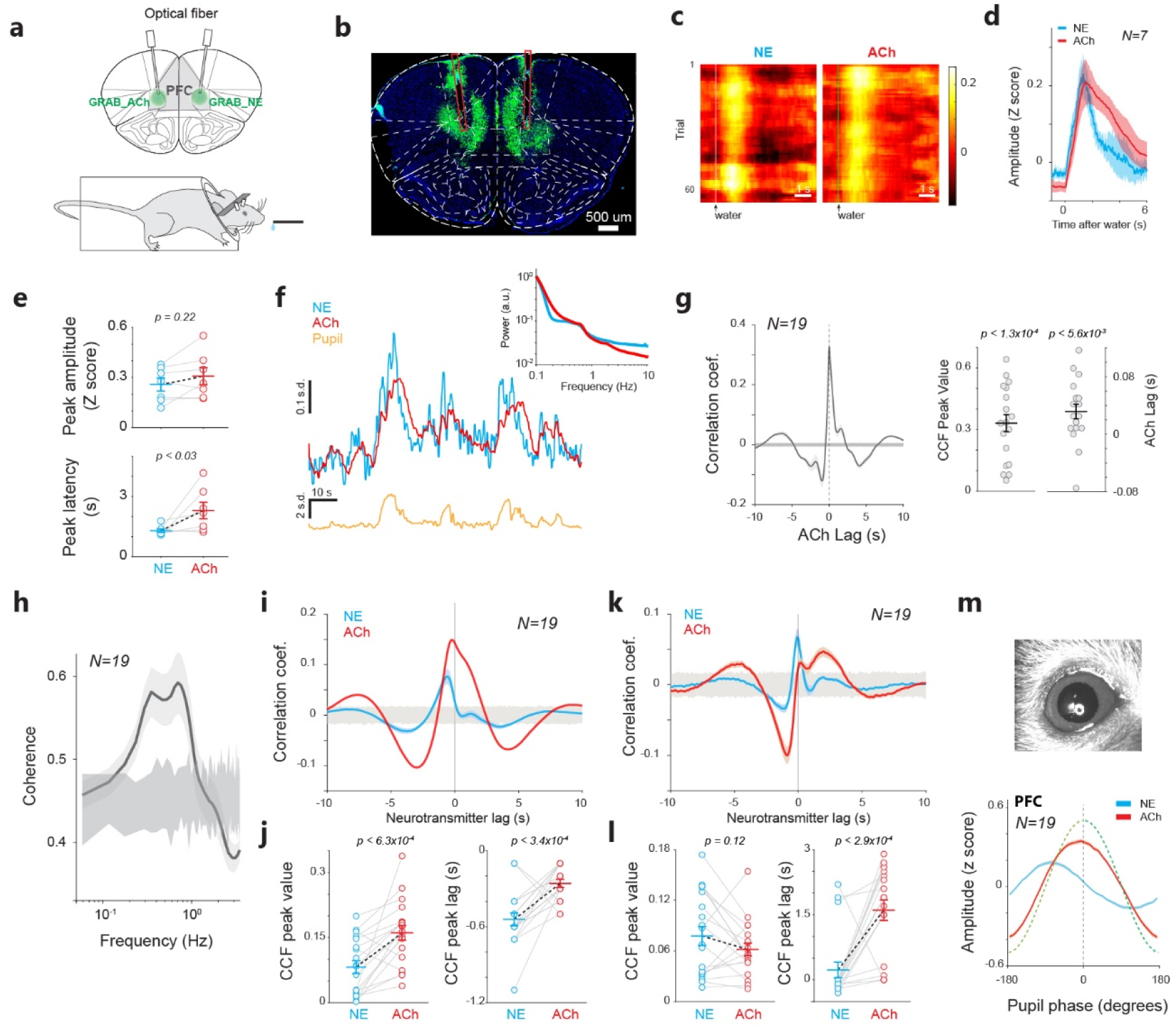
Spontaneous fluctuation of NE and ACh levels in the prefrontal cortex. **a)** Diagram of GRAB_NE_ and GRAB_ACh_ recording. **b)** Histological confirmation of expression of GRAB_NE_ and GRAB_ACh_ in the prefrontal cortex. **c)** Example heatmap of NE and ACh responses to water rewards. **d)** NE and ACh dynamics around water rewards. 25 sessions from 7 animals. **e)** NE and ACh peak responses (top) and their latency (bottom) to water rewards. **f)** Example traces of simultaneously recorded NE, ACh, and pupil size. Inset: spectrum of NE and ACh signals. **g)** Cross-correlogram between NE and ACh signals. 165 sessions from 19 animals. Shaded area around 0 indicates 99.7% confidence interval. **h)** Coherence between NE and ACh signals. 165 sessions from 19 animals. Shaded area around 0 indicates 99.7% confidence interval. **i,j)** Cross-correlogram between NE/ACh signals and pupil size. 111 sessions from 19 animals. Horizontal shaded area around 0 indicates 99.7% confidence interval. **k,l)** Cross-correlogram between NE/ACh signals and the first derivative of pupil size. 111 sessions from 19 animals for panels k-m. Shaded area around 0 indicates 99.7% confidence interval. **m)** Example image of the pupil of a mouse (top) and the phase relationship between prefrontal NE/ACh signals and pupil fluctuations (bottom). Error bars or shaded area indicate S.E.M. in all figures unless otherwise indicated.

Prior studies have established that pupil size can track the activity of noradrenergic and cholinergic axons in the sensory cortices ^32^. To further examine if pupil size can also track NE and ACh fluctuations in the prefrontal cortex, we performed cross-correlation analysis between pupil size and GRAB_NE_/GRAB_ACh_ signals. As expected, the cross-correlogram revealed a positive correlation between pupil size and NE/ACh signals, with ACh exhibiting a slightly higher peak correlation coefficient (p<6.3e-4; paired t-test) (**Figure 1i, j**). We also found that both NE and ACh signals preceded pupil fluctuations (NE: -0.54±0.05 s, significantly differs from 0 with p<1.2e-4, Wilcoxon signed-rank test; ACh: -0.26±0.03 s, significantly differs from 0, p=1.2e-4, Wilcoxon signed-rank test). Consistent with the direct cross-correlation results between NE and ACh, NE signals preceded pupil signals more than ACh signals did (p<3.4e-4, paired t-test) (**Figure 1j**). As tonic LC stimulation evoked continuous pupil dilation ^33^, we further correlated NE/ACh signals with the first derivative of pupil size ^32^ (**Figure 1k**). Interestingly, although the peak value of the cross-correlogram between NE signals and pupil size derivative was comparable to that between ACh signals and pupil derivative (p=0.12, paired t-test), the peak latency was larger for ACh than NE signals (p<2.9e-4, paired t-test, **Figure 1l**). The correlated fluctuations between cortical NE/ACh signals and pupil size were further confirmed by aligning NE and ACh fluctuations to one canonical cycle of pupil dilation and constriction derived from the Hilbert transform. Similar to previous findings^32^, we found that both NE and ACh exhibited peak amplitude at a negative pupil phase, confirming that both NE and ACh signals preceded pupil fluctuations (**Figure 1m; Supplemental figure 1b,c**).

#### Prefrontal NE and ACh dynamics during inhibitory control

After verifying the functionality of the biosensors, we focused on understanding the NE and ACh dynamics during inhibitory control. In pursuit of this, we simultaneously measured the NE and ACh signals in the prefrontal cortex from 19 mice during an inhibitory control task (**Figure 2a**). In this task, mice were required to withhold their impulsive licking. During the initial shaping period, once naïve mice associated the waterspout with sweet water delivery (usually on the first day), they constantly licked to check for sweet water even though a sweet water drop was randomly delivered every 12-22 seconds. This habitual behavior was evidenced by elevated licking frequencies during subsequent sessions (**Figure 2b**, p<8.1e-6, one-way ANOVA test). In the inhibitory control task, mice were trained to suppress the impulsive licking. At the beginning of each trial, the animals could freely lick with no penalty during a free period (5 to 7 s uniform distribution). Subsequently, an inhibition tone (duration randomly drawn from an exponential distribution varying from 5 to 12 s with λ=4.5, **Figure 2a**) was played. During the inhibition tone period, a lick would trigger a brief air puff (20 psi, 200 ms, see **Methods**) to the animal’s face and immediately terminate the inhibition tone, while successful withholding of licking would result in a sweet water reward at the end of the inhibition tone. Our data demonstrated that the mice could effectively suppress their impulsive licking once the tone started, and their success rate gradually increased during the initial training sessions, suggesting that their inhibitory control is a learned behavior (**Supplemental figure 2a**). The licking frequency within 2 seconds following the inhibition tone onset was significantly lower than during a 2-second window immediately prior to the tone onset (**Figure 2c, Supplemental figure 2b,** p<4.2e-10; Wilcoxon signed-rank test). Furthermore, the animals’ success rates were significantly higher than the chance levels (**Figure 2d**, p<3e-10, paired t-test; **Supplemental figure 2c**). As we expected, the longer the inhibition tone period (trials were grouped into three inhibition tone periods: 5-7.5, 7.5-10, and 10-12 seconds), the less likely that the mice were able to suppress their impulsive licking (**Figure 2e**, p<7.4e-10, one-way ANOVA test). The animals typically collected water rewards within 500 ms (**Supplemental figure 2d**). The duration of the inhibition tone had no effect on the reaction time, defined as the interval between the offset of the inhibition tone and the animal’s first lick to collect water reward, suggesting that the animals were generally vigilant during the task (**Figure 2f**, p=0.96, one-way ANOVA test). Together, these behavioral results suggested that the animals exercised cognitive control to suppress impulsive licking in the inhibitory control task.

**Figure 2.**
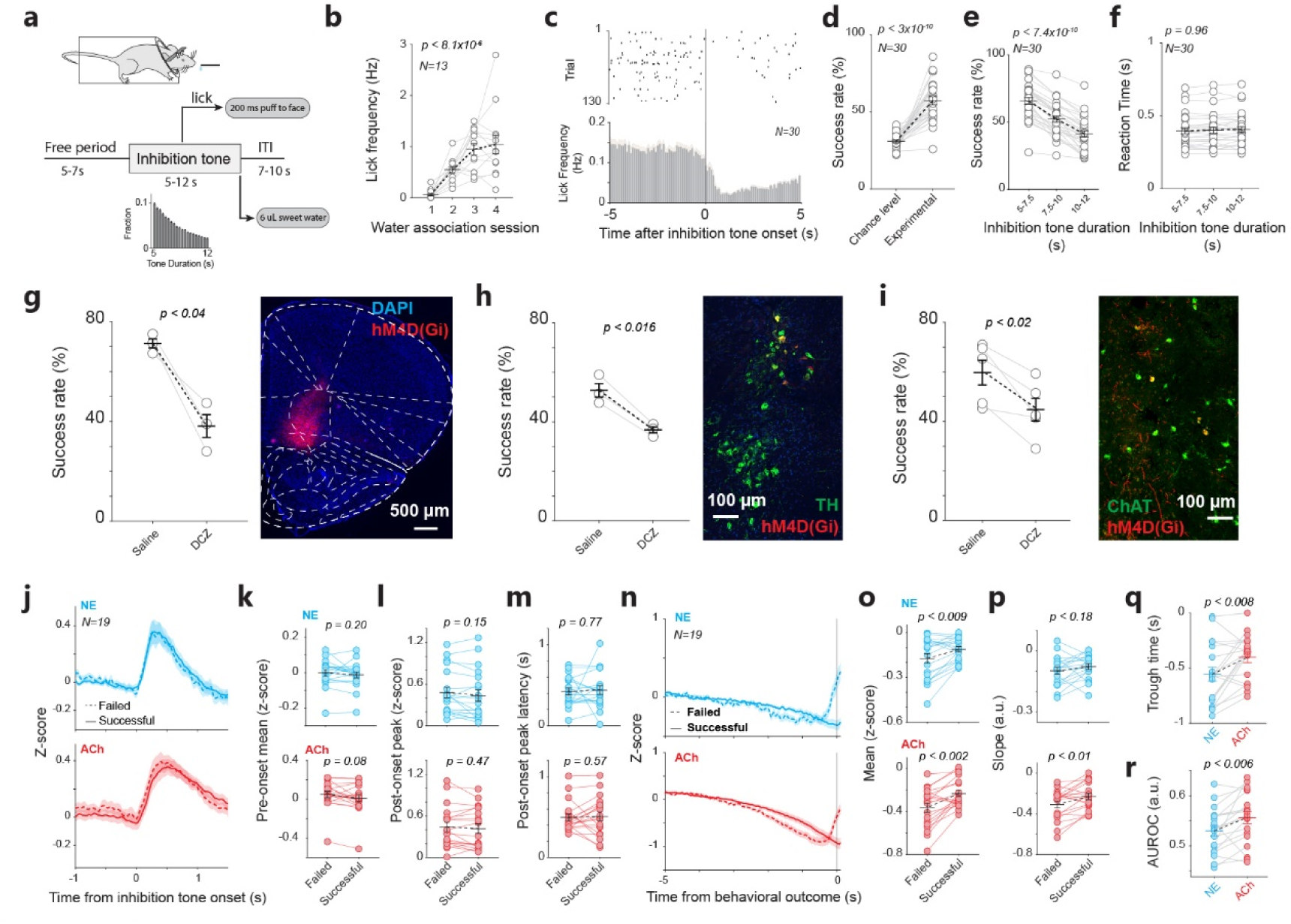
Prefrontal NE and ACh dynamics during inhibitory control. **a)** Diagram of the inhibitory control task. **b)** Impulsive licking frequency during the initial shaping period. 52 sessions from 13 animals. **c)** Example raster plot of licks (top) and average licking frequency (bottom) around the onset of the inhibition tone. 260 sessions from 30 animals for panels c-f. **d)** Raw success rate and the chance-level success rate. **e)** Raw success rate associated with different inhibition tone durations. **f)** Reaction time associated with different inhibition tone durations. **g)** Success rate with and without the inactivation of the prefrontal cortex. 28 sessions from 3 animals. **h)** Success rate with and without the inactivation of noradrenergic inputs to the prefrontal cortex. 30 sessions from 3 animals. **i)** Success rate with and without the inactivation of cholinergic inputs to the prefrontal cortex. 44 sessions from 5 animals. **j)** NE and ACh dynamics around the onset of inhibition tone for the successful and failed trials. 165 sessions from 19 animals for panels g-o. **k)** Mean NE and ACh levels before inhibition tone onset. **l)** The peak value of NE and ACh transient responses to inhibition tone onset. **m)** The peak latency of NE and ACh transient responses. **n)** NE and ACh dynamics prior to behavioral outcomes in the successful and failed trials. **o)** Mean NE and ACh levels prior to behavioral outcomes. **p)** The slope of NE and ACh signals prior to behavioral outcomes. **q)** The trough time of NE and ACh signals prior to behavioral outcomes. **r)** The area under ROC curve (AUROC) calculated from signal distributions associated with the successful and failed trials for NE and ACh signals

Although previous work using pharmacological manipulations suggested an important role of NE in the PFC in inhibitory control ^34^, we first examined whether the PFC is required for the task used in our study. Chemogenetic inactivation of the PFC in 3 WT mice significantly impaired their success rate in the inhibitory control task (**Figure 2g**, p<0.04, paired t-test; **Supplemental figure 3a**), suggesting that the PFC is necessary for the inhibitory control task. We next examined whether noradrenergic or cholinergic signals in the PFC are required for inhibitory control. Chemogenetic inactivation of noradrenergic inputs to the PFC resulted in a reduction of 15.9% in success rate (**Figure 2h**, p<0.016, paired t-test; **Supplemental figure 3b**), while chemogenetic inactivation of cholinergic inputs to the PFC decreased animals’ success rate by 14.9% (**Figure 2i**, p<0.02, paired t-test; **Supplemental figure 3c**). Once we confirmed the necessity of NE and ACh signals in the PFC for the inhibitory control task, we started to explore prefrontal NE and ACh dynamics during inhibitory control.

Consistent with previous work demonstrating phasic LC firing in response to salient stimuli ^21,25,35,36^, as we expected, the onset of inhibition tone elicited a phasic increase of NE levels in the PFC (**Figure 2j)**. Interestingly, ACh concentration also exhibited a dramatic increase following the onset of inhibition tone. When comparing the phasic responses between successful and failed trials, we found that there was no significant difference in either NE or ACh levels prior to the inhibition tone between the two behavioral outcomes (**Figure 2k**, NE: p=0.20, Wilcoxon signed-rank test; ACh: p=0.08, Wilcoxon signed-rank test). We also failed to find significant differences in the peak amplitude or latency of evoked transient NE and ACh responses between successful and failed trials (**Figure 2l,m**; peak amplitude: NE: p=0.15, Wilcoxon signed-rank test; ACh: p=0.47, Wilcoxon signed-rank test. Latency: NE: p=0.77, Wilcoxon signed-rank test; ACh: p=0.57, Wilcoxon signed-rank test).

Given our results suggesting that NE and ACh dynamics prior to the inhibition tone do not index inhibitory control performance, we then examined the NE and ACh dynamics in the PFC within the 5 seconds preceding the behavioral outcome for each trial (i.e. reward at the end of inhibition tone or punishment resulting from licking during the inhibition tone). We chose the 5- second window because it represents the minimum duration of the inhibition tone so we can include all successful trials in our analysis. Although both NE and ACh signals were generally decreasing before both behavioral outcomes, they initiated an increase at approximately 0.5 s before impulsive licking in failed trials (**Figure 2n-p)**. We found that there was a difference in the extracellular level of both neurotransmitters between successful and failed trials (**Figure 2o**; NE: p<0.009, Wilcoxon signed-rank test; ACh: p<0.002, Wilcoxon signed-rank test). The difference in the descending slope of NE/ACh signals was only significant for ACh, not for NE (**Figure 2p**; NE: p=0.18, Wilcoxon signed-rank test; ACh: p<0.01, Wilcoxon signed-rank test). The trough time for NE slightly preceded ACh (**Figure 2q**, p<0.008, paired t-test). To further quantify the discriminability between NE/ACh levels in successful versus failed trials, we performed ROC analysis ^37^. In this analysis, the area under the ROC curve (AUROC) is a quantitative measure of the discriminability (i.e., normalized difference) between two stochastic signals. Consistent with the results shown in **Figure 2n**, the AUROC associated with ACh signals was greater than that associated with NE signals, suggesting that ACh dynamics associated with successful and failed outcomes were more separated than NE (**Figure 2r**, p<0.006, Wilcoxon signed-rank test).

We also performed a support vector machine (SVM) classifier analysis to evaluate the discriminability of single-trial NE/ACh dynamics between successful and failed trials. Consistent with the AUROC analysis, the classifier performed better at classifying single-trial ACh dynamics associated with successful or failed trials than NE dynamics (**Supplemental figure 4a**, p < 0.01, paired t-test). Furthermore, we did not observe a significant difference in the correlation between NE and ACh dynamics during inhibitory control across successful and failed trials (**Supplemental figure 4b**, p=0.28, paired t-test).

#### The dynamics of either NE or ACh in the PFC are not reliable indicators of inhibitory control

Our results just suggest that both NE and ACh levels in the prefrontal cortex appear to be linked to inhibitory control, as the neurotransmitter levels associated with success or failure in impulse control differed for both NE and ACh. However, if NE/ACh levels prior to behavioral outcomes truly index inhibitory control, their difference between successful and failed trials would diminish or even vanish if inhibitory control is impaired. To test this, we conducted experiments in which we manipulated the LC-NE system in the behaving mice. To this end, we injected AAVrg-hSyn-DIO-hM4D(Gi)-mCherry into the basal forebrain region of DBh-Cre mice to retrogradely express inhibitory DREADD receptors in LC neurons that project to the basal forebrain area (**Figure 3a**, see **Methods**). Post-mortem IHC confirmed the expression of DREADD in LC neurons (**Figure 3b** and **Supplemental figure 5a**). In control sessions where the mice received saline administration, their inhibitory control performance was significantly greater than the chance level (**Supplemental figure 5b,** p<0.02, paired t-test), similar to WT mice (**Supplemental figure 5c, d**). However, when CNO was administered to inactivate LC neurons projecting to the basal forebrain region, the mice’s performance dramatically dropped by 19.6% (**Figure 3c**. p<4e- 3, paired t-test; **Supplemental figure 5b**). Indeed, the performance with CNO was not significantly different from the chance level (p=0.28, paired t-test), suggesting their inhibitory control was totally impaired. To control for the effect of CNO alone (i.e., not through its effect on DREADD receptors), we administered CNO/DCZ to 4 WT mice, and found that CNO/DCZ alone had no effect on these animals’ inhibitory control (**Supplemental figure 6a)**.

**Figure 3.**
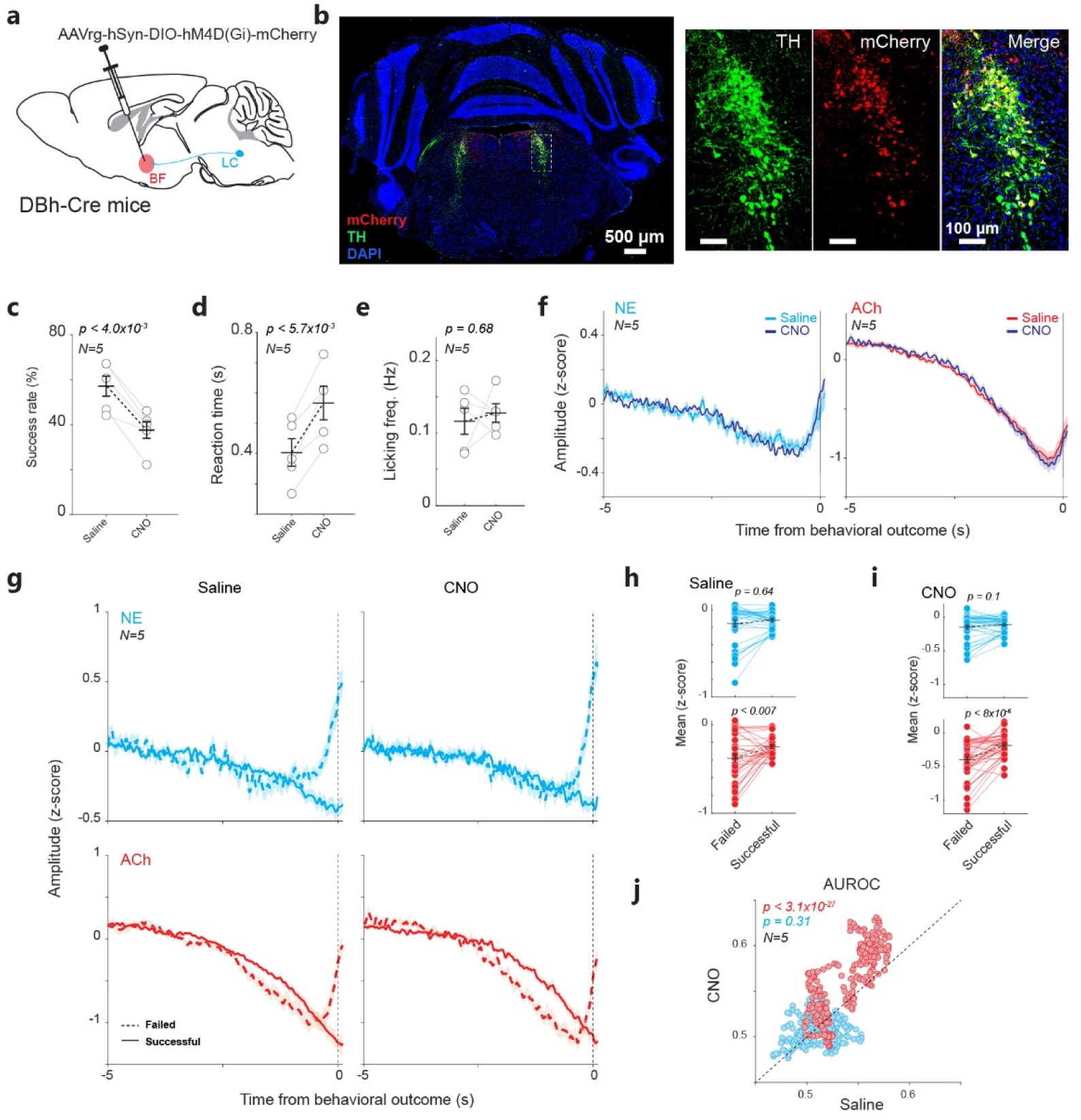
Chemogenetic silencing of BF-projecting LC neurons impaired the behavior but did not diminish the difference in NE and ACh signals between successful and failed trials. **a)** Diagram of retrograde expression of DREADD receptors in LC neurons that project to the basal forebrain region. **b)** Histological confirmation of the expression of DREADD receptors in LC neurons. **c)** CNO-mediated chemogenetic inhibition of LC neurons that project to the basal forebrain region reduced the inhibitory control performance to the chance level. **d,e)** Chemogenetic inhibition of LC neurons that project to the basal forebrain region slowed down the reaction time but did not change licking frequency during the free period. **f)** Average NE/ACh signals prior to behavioral outcomes under saline and CNO treatment. **g)** NE/ACh signals prior to behavioral outcomes in the successful and failed trials under saline and CNO treatment. **h,i)** Mean NE and ACh levels prior to behavioral outcomes in the successful and failed trials under saline and CNO treatment. **j)** Area under the ROC curve (AUROC), which measures the normalized difference in NE/ACh levels between the successful and failed trials, under saline and CNO treatment. All data are from 34 saline sessions and 42 CNO sessions from 5 animals.

While CNO administration slowed their reaction time (**Figure 3d**, p<5.7e-3, paired t-test), it did not significantly affect the licking frequency during the free period (**Figure 3e**, p=0.68, paired t- test). Interestingly, there was an inconspicuous difference in NE/ACh dynamics before behavior outcomes between CNO treatment sessions and saline control sessions (**Figure 3f**). CNO-mediated inhibition of LC neurons projecting to the basal forebrain region also mildly affected prefrontal NE and ACh dynamics induced by the inhibition tone or water reward (**Supplemental figure 6b,c**). Because CNO-mediated inhibition of LC neurons greatly disrupted inhibitory control, we expected it also to abolish the difference in NE/ACh levels between successful and failed trials. Surprisingly, when we segregated the NE and ACh signals based on behavioral outcomes, contrary to what we expected, we found that the inactivation of LC neurons did not diminish the difference between successful and failed trials for either NE or ACh signals. Instead, the difference in the ACh signal appeared to be enhanced while the inhibitory control performance was at the chance level, suggesting that the difference in mean NE/ACh signals between successful or failed trials is not a reliable indicator of inhibitory control (**Figure 3g-i**). To quantify these results, we again used ROC analysis. The AUROC from ACh signals indeed increased, while AUROC from NE signals remained the same (**Figure 3j**. ACh: p<3.1e-27, Wilcoxon signed-rank test; NE: p=0.31, Wilcoxon signed-rank test). Taken together, these results suggest that the averaged NE or ACh signals are not correlated with inhibitory control. SVM analyses also yielded slightly higher classification accuracy for ACh signals when LC neurons projecting to the basal forebrain region were inhibited. However, this difference was not statistically significant (**Supplemental figure 4c**).

#### Phase synchrony between NE and ACh signals in the PFC indexes inhibitory control

Given that our results suggested that mean NE and ACh levels are not reliably linked to inhibitory control, we then asked what other features of the NE and ACh signals could be reliably linked to the behavior. As the NE and ACh system exhibited the strongest coherence at 0.4-0.8 Hz, we first explored whether the phase of NE and ACh fluctuations at this frequency band depends on inhibitory control (**Figure 4a**). We found that CNO-induced inhibition of LC neurons projecting to the basal forebrain did not eliminate the difference in the phase distributions of ACh and NE signals between successful and failed trials, suggesting the phase is also not linked to inhibitory control (**Figure 4b**). Upon closer examination of NE and ACh signal fluctuations, we noticed that the phase relationship between NE and ACh signals was dynamic. Specifically, NE and ACh exhibited similar phases (i.e. in-phase) during some periods but had opposite phases (i.e. out-of-phase) during the other periods. This phenomenon has been observed in other physiological signals ^38^. We calculated the value of a phase encoder as a quantitative measure of phase synchrony between NE and ACh signals (see **Methods**) ^38^. In this measure, a value of one indicates that the two signals share the same phase angle and thus are in-phase, while the value of zero indicates that the two signals have a phase difference of 180 degrees (i.e. out-of-phase). CNO-mediated inhibition of LC neurons projecting to the basal forebrain did not significantly alter the overall phase synchrony during the 5 second period before behavioral outcomes (**Figure 4c**). However, while the phase synchrony between successful and failed trials began to diverge approximately 3 seconds before behavioral outcomes in saline control sessions (**Figure 4d, left panel**), the inhibition abolished the difference in phase synchrony between successful and failed trials (**Figure 4d, right panel; Figure 4e**, Saline: p<0.004, paired t-test; CNO: p=0.26, paired t-test), resulting in a decreased AUROC with CNO compared to saline controls (**Figure 4f**, p<2.6e-36, Wilcoxon signed-rank test). Because ACh signals lag behind NE signals by 30 ms, we performed a computational control to ensure that the observed difference in phase synchrony between successful and failed trials during inhibitory control was not an artifact of this lag. To this end, we shifted ACh signals 30 ms forward and used this shifted ACh signals and the original NE signals to calculate phase synchrony in successful and failed trials. We still observed a substantial difference in phase synchrony between successful and failed trials, which disappeared when LC neurons projecting to the basal forebrain were inhibited. (**Supplemental figure 7a**).

**Figure 4.**
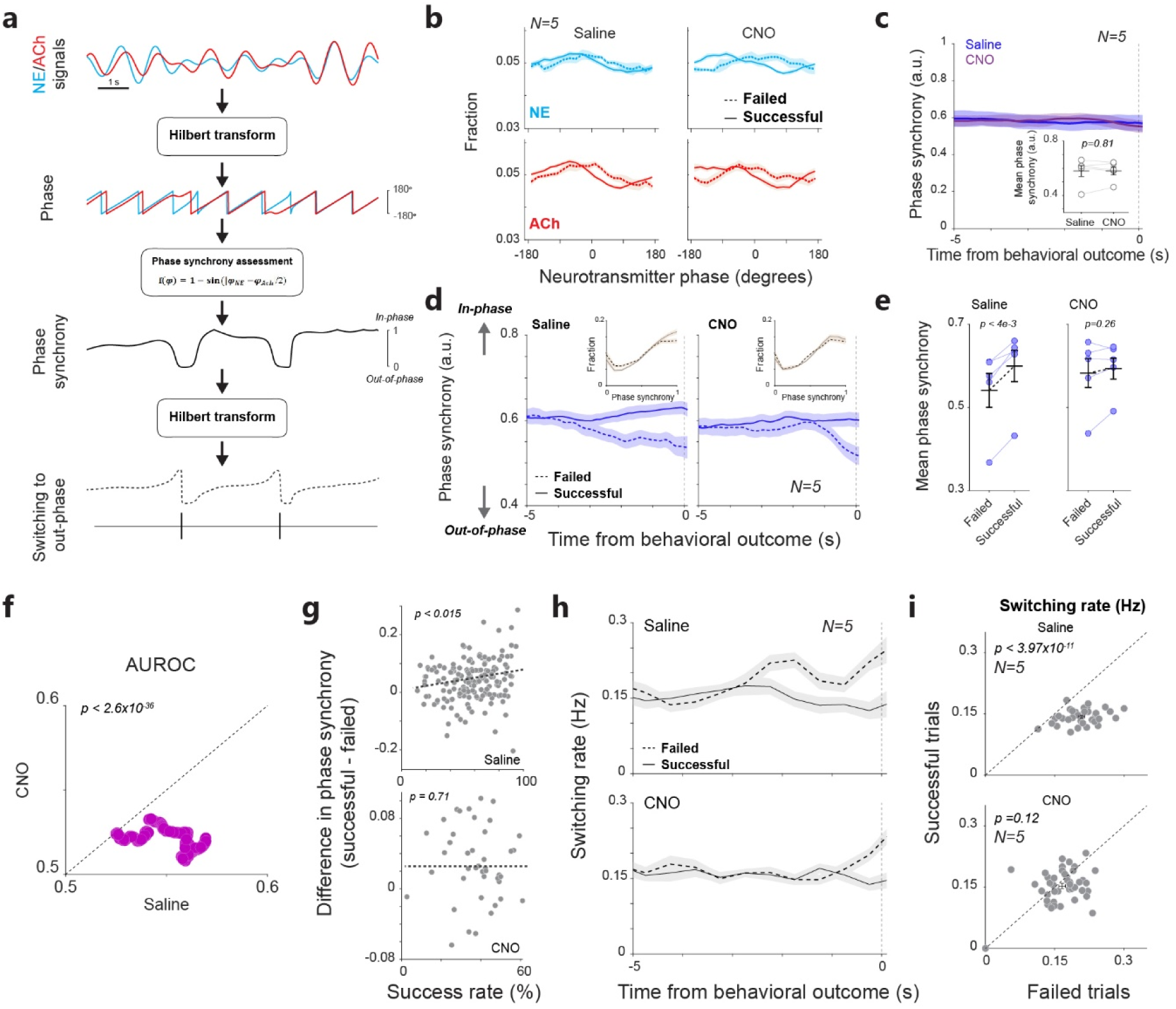
Phase synchrony between prefrontal NE and ACh signals is a robust indicator of inhibitory control behavior. **a)** Illustration of the estimation of NE-ACh phase synchrony. **b)** Distribution of NE/ACh phase in successful and failed trials under saline and CNO treatment. 34 saline sessions and 42 CNO sessions from 5 animals for panels c-h. **c)** NE-ACh phase synchrony prior to behavioral outcomes under saline and CNO treatment. **d)** NE-ACh phase synchrony prior to behavioral outcomes in successful and failed trials under saline and CNO treatment. **e)** Mean NE-ACh phase synchrony prior to behavioral outcomes in successful and failed trials under saline and CNO treatment. **f)** AUROC calculated using NE-ACh phase synchrony under saline vs. CNO treatment. **g)** Difference in prefrontal NE-ACh phase synchrony between successful and failed trials was positively correlated with inhibitory control performance in saline control sessions but not in CNO treatment sessions. **h)** Switching rate prior to behavioral outcomes in successful and failed trials under saline and CNO treatment. **i)** Mean switching rate prior to behavioral outcomes in successful vs. failed trials under saline or CNO treatment. 76 sessions from 5 animals.

Furthermore, the difference in NE-ACh phase synchrony between successful and failed trials was positively correlated with behavioral performance and this correlation again vanished when BF-projecting LC neurons were inhibited by CNO (**Figure 4g, Supplemental figure 7b**), suggesting that the NE and ACh interplay in the prefrontal cortex is reliably linked to inhibitory control. While the average phase synchrony between NE and ACh signals in the PFC was generally weaker in failed trials than in successful trials, we wondered whether there were other features of the phase synchrony linked to inhibitory control. We found that the distribution of phase synchrony was non-uniform. Rather, the distribution was heavily skewed towards the value of 1, indicating that the NE and ACh activities were mostly in phase (**Figure 4d**, insets). Indeed, upon close inspection of the phase synchrony, we observed occasional rapid decreases to around 0, signifying a transient out-of-phase state between the noradrenergic and cholinergic systems for a short period (**Figure 4a**, second to the bottom panel). Although the mechanism through which the NE and ACh systems were transiently out-of-phase remains unknown, we examined whether switching from in-phase to out-of-phase between the NE and ACh systems was linked to behavior. To this end, we assessed the number of switches before the behavioral outcome. We found a noticeable difference in the switching rate between successful and failed trials approximately 3 seconds before the behavioral outcome (**Figure 4h**). More importantly, similar to its effect on behavioral performance, the CNO-mediated silencing of LC neurons projecting to the basal forebrain region significantly reduced the difference between successful and failed outcomes (**Figure 4i**. Saline: p<3.98e-11, paired t-test; CNO: p=0.12, paired t-test). For all 19 animals in which we recorded NE and ACh from the prefrontal cortex, the difference in the switching rate between successful and failed trials appeared to be positively correlated with their behavioral performance (**Supplemental figure 7c,** p<0.005).

#### Inactivation of cholinergic projections to the LC did not affect prefrontal NE-ACh phase synchrony and inhibitory control

As silencing LC neurons projecting to the basal forebrain regions profoundly disrupted inhibitory control and behavior-dependent phase synchrony, we further explored if possible basal forebrain projections to the LC have a similar effect on inhibitory control ^39,40^. We again injected the retrograde AAV virus into the LC of ChAT-Cre mice to retrogradely express inhibitory DREADD receptors in cholinergic neurons projecting to the LC (**Supplemental figure 8a**). IHC analysis demonstrated the expression of mCherry fluorescent protein, the tag of the retrograde AAV, in the superior cerebellar peduncle (SCP)/Red nucleus (RN), the pontine reticular nucleus (PRN), and the trigeminal motor nucleus (V) area (**Supplemental figure 8b**). Interestingly, we did not find evident mCherry expression in the basal forebrain area in these animals (see **Discussion**). Silencing the cholinergic projections to the LC failed to disrupt the inhibitory control performance (**Supplemental figure 8c**, p=0.55, paired t-test). However, the reaction time for the animals to collect water rewards at the offset of the inhibition tone in successful trials was significantly slower than in saline control sessions (**Supplemental figure 8d**, p<0.001, paired t-test), indicating the effects of the chemogenetic inhibition of cholinergic neurons on motor-related functions, but not on cognitive functions.

Similar to the results from manipulating the LC-NE system in the DBh-Cre mice, manipulating the cholinergic projections to the LC failed to diminish the differences in prefrontal NE and ACh signals between successful and failed trials (**Supplemental figure 8e**). Consistent with their effect on behavioral performance, silencing the cholinergic projections to the LC failed to modulate the phase synchrony between the NE and ACh signals (**Supplemental figure 8f**). Neither did it significantly change the switching rate as there was a higher switching rate in failed trials than in successful trials for both CNO treatment and the saline controls (**Supplemental figure 8g**, Saline: p<3.1e-6, paired t-test; CNO: p<4.7e-6, paired t-test). Taken together, these results strengthened the notion that prefrontal NE-ACh phase synchrony is linked to inhibitory control.

#### Distinct subgroups of LC neurons that project to the PFC and BF

Our data demonstrate that inhibiting LC neurons projecting to the basal forebrain significantly impaired inhibitory control behavior and disrupted phase synchrony between prefrontal NE and ACh signals, while having only a limited impact on prefrontal NE amplitudes. One possibility is that LC neurons projecting to the basal forebrain are distinct from those projecting to the PFC ^41–43^. To test this hypothesis, we conducted AAV-mediated Cre-dependent dual-color retrograde tracing in DBh-Cre mice. We simultaneously injected AAVrg-EF1a-DIO-hChR2(H134R)-mCherry (or AAVrg-EF1a-DIO-hChR2(H134R)-EYFP) into the orbitofrontal region (OFC) of the PFC, and AAVrg-EF1a-DIO-hChR2(H134R)-EYFP (or AAVrg-EF1a-DIO-hChR2(H134R)-mCherry) into the basal forebrain in 4 DBh-Cre mice (**Figure 5a**. See **Methods**). We used these two AAV vectors as retrograde tracers because they share the same construct, differing only in their fluorescent tags. Approximately 3–4 weeks after injection, we sectioned the brain throughout the LC and used immunohistology to quantify the number of LC neurons expressing mCherry, EYFP, or both (**Figure 5b,c**). Along the anterior-posterior axis, we found that OFC-projecting LC neurons are distributed throughout the LC, but with their highest density located approximately 250 µm from the start of the LC (**Figure 5d**). In contrast, BF-projecting LC neurons are relatively evenly distributed along the anterior-posterior axis of the LC (**Figure 5d**). The similar distribution pattern was found for LC neurons that project to both the PFC and BF (**Figure 5d**).

**Figure 5.**
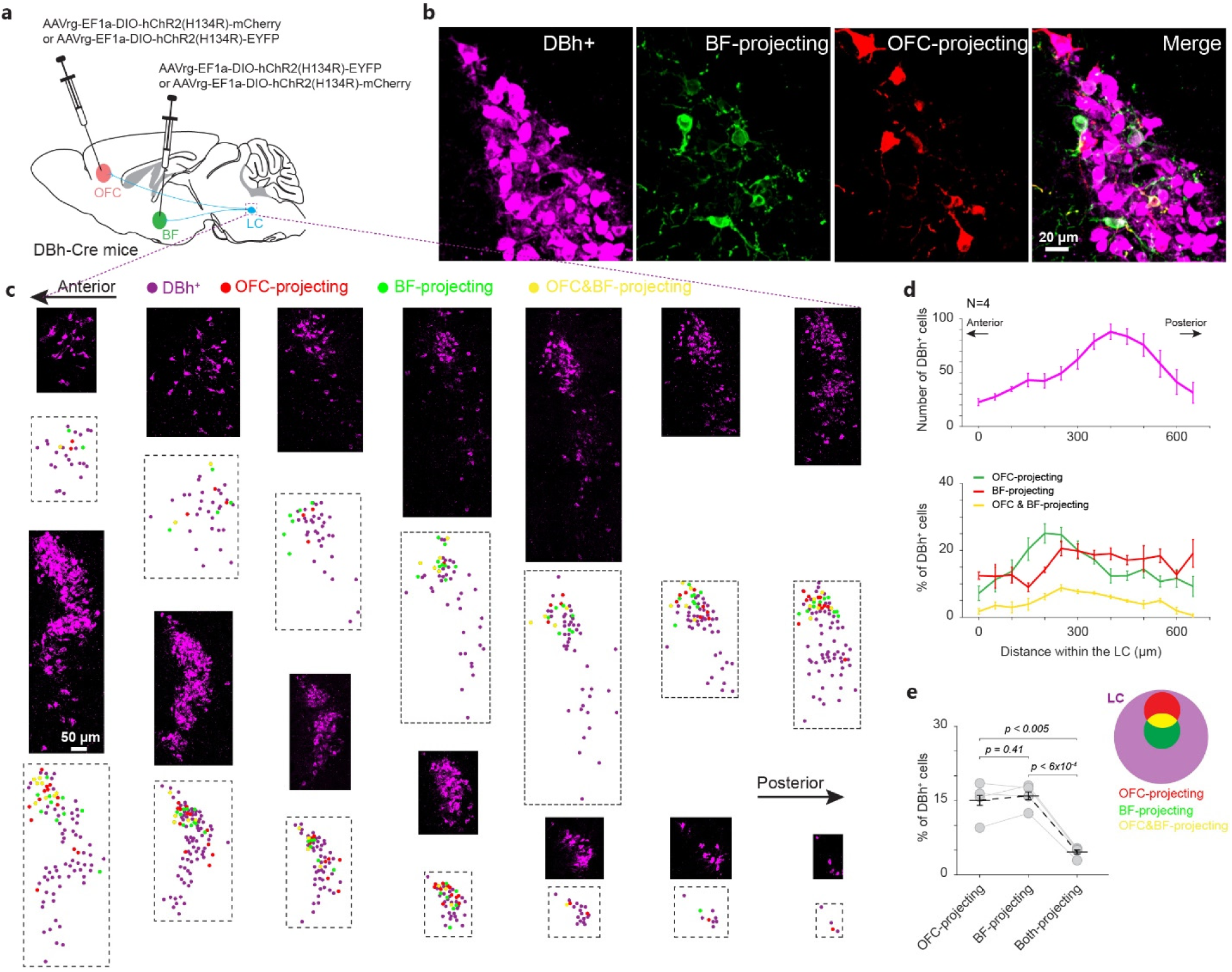
Retrograde tracing revealed distinct subgroups of LC neurons projecting to the prefrontal cortex and basal forebrain. **a)** Diagram showing retrograde expression of different fluorophores in LC neurons that project to the prefrontal cortex and basal forebrain. **b)** Example confocal image showing co-expression of EYFP (pseudocolored green) and mCherry in LC neurons. **c)** Sections of the LC from an example mouse illustrating the spatial distribution of OFC-projecting and BF-projecting LC neurons. **d)** Quantification of LC neurons, OFC-projecting, and BF-projecting LC neurons across anterior-posterior sections. **e)** Overall quantification of LC neurons projecting to the OFC, BF, or both. The cartoon illustrates the percentage of LC neurons projecting to the PFC, BF, or both.

In each LC section, OFC-projecting and BF-projecting LC neurons are primarily located in the dorsal part of the LC. These two subgroups are intermingled, forming a salt-and-pepper pattern (**Figure 5c**). Overall, the number of OFC-projecting LC neurons is comparable to the number of BF-projecting LC neurons (**Figure 5e**, p=0.41, paired t-test), each accounting for approximately 15% of LC neurons. LC neurons projecting to both the PFC and BF constitute only about 30% of OFC-projecting or BF-projecting LC neurons (**Figure 5e**).

#### Prefrontal population activity during inhibitory control

Previous work has demonstrated the critical role of the prefrontal cortex in inhibitory control. After discovering that prefrontal NE-ACh phase synchrony is a behaviorally relevant neuromodulatory feature, we further investigated its potential effect on the prefrontal neural activity that mediates inhibitory control. We implanted a Neuropixels probe into the prefrontal cortex to record population activity while chemogenetically inhibiting LC neurons that project to the basal forebrain during inhibitory control (**Figure 6a**). Active neurons (firing rate > 1 Hz; see **Methods**) during inhibitory control were located in various regions of the prefrontal cortex, including the orbitofrontal and prelimbic regions. This suggested that these sub-regions of the prefrontal cortex contribute to inhibitory control. 17% of the neurons exhibited a narrow waveform and were therefore termed fast-spiking units (FSUs), while the remainder were termed regular-spiking units (RSUs) due to their broader waveforms (**Figure 6b**). Both RSUs and FSUs were evenly distributed along the probes over a distance of approximately 1.1 mm from the electrode tip. Consistent with results from the biosensor group, the inhibition of BF-projecting LC neurons significantly impaired the animal’s inhibitory control performance (**Figure 6c**, p<0.015, paired t test). Moreover, the firing rate of prefrontal neurons during inhibitory control was decreased during DCZ sessions, confirming LC modulation on prefrontal activity (**Figure 6d**, p<0.035, t-test). DCZ-mediated inhibition of LC neurons projecting to the basal forebrain also had a mild impact on prefrontal population firing rate around the onset of the inhibition tone (**Supplemental figure 9**).

**Figure 6.**
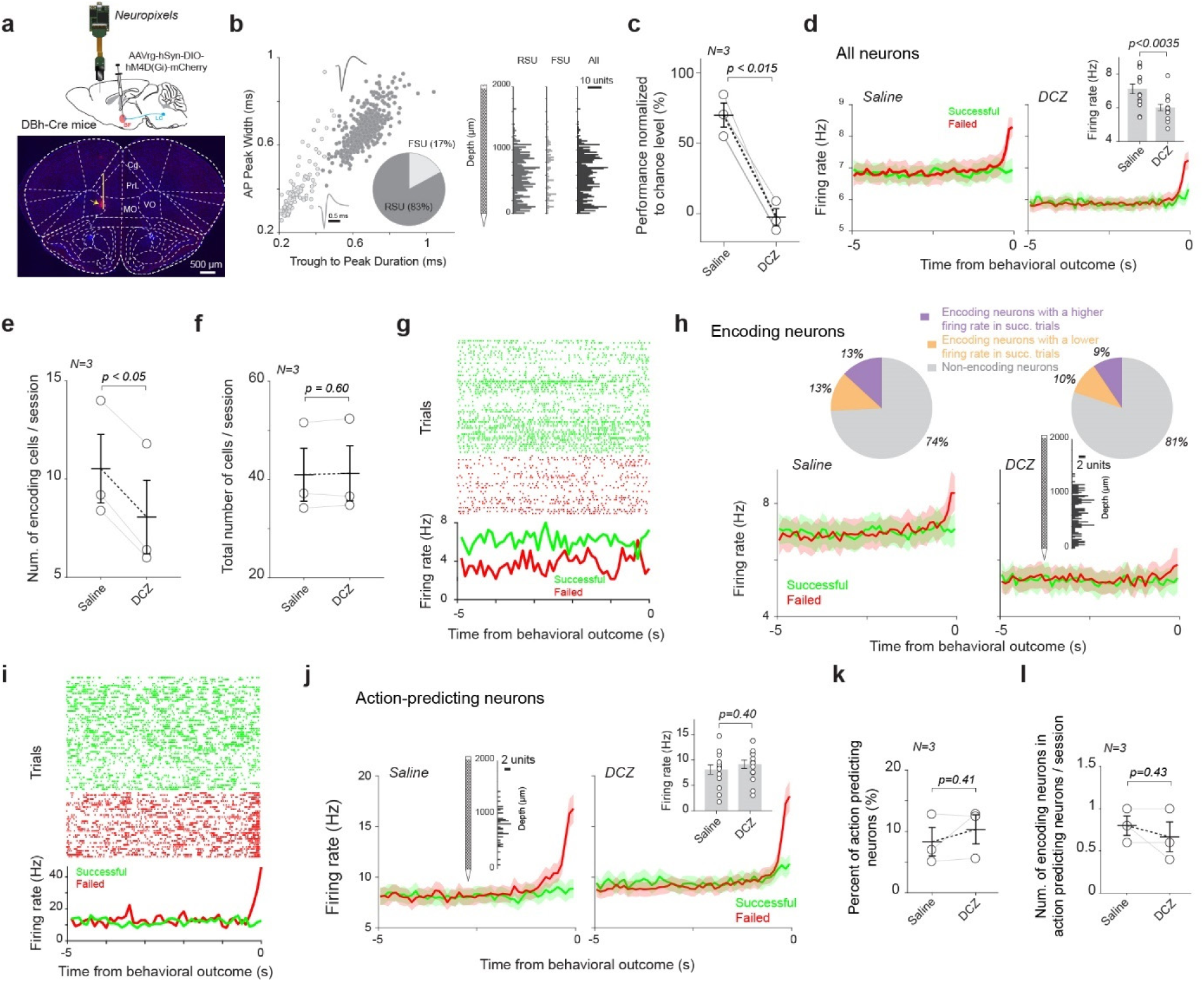
Neuropixels recording from the prefrontal cortex during inhibitory control. **a)** Histological confirmation of the location of the Neuropixels probe (indicated by red dye and the yellow arrow) in the prefrontal cortex. The yellow line indicates a distance of 1.2 mm from the Neuropixels probe tip, where most of the active units during inhibitory control were located. **b)** Waveform characteristics of regular spiking units (RSU) and fast spiking units (FSU) (left panel) and their location from the tip of the Neuropixels probe (right panel). 15 saline sessions and 15 DCZ sessions from 3 animals for all neuropixels results. **c)** DCZ-mediated chemogenetic inhibition of LC neurons that project to the basal forebrain region reduced the inhibitory control performance to the chance level. **d)** Population firing rate prior to behavioral outcomes in the successful and failed trials under saline and DCZ treatment. Inset: mean firing rate under saline and DCZ treatment. **e)** Number of encoding neurons under saline and DCZ treatment. **f)** Total number of neurons under saline and DCZ treatment. **g)** Raster plot of spikes of an example encoding neuron on successful and failed trials. **h)** Population firing rate of encoding neurons prior to behavioral outcomes in the successful and failed trials under saline and DCZ treatment. Pie charts: percentage of encoding neurons with higher firing rates in successful trials and encoding neurons with lower firing rates in successful trials. Inset: distribution of encoding neurons along neuropixels probe. **i)** Raster plot of spikes of an example action-predicting neuron on successful and failed trials. **j)** Population firing rate of action-predicting neurons prior to behavioral outcomes in the successful and failed trials under saline and DCZ treatment. Left inset: distribution of encoding neurons along neuropixels probe. Right inset: mean firing rate under saline and DCZ treatment. **k)** Percentage of action-predicting neurons under saline and DCZ treatment. **l)** Number of encoding neurons among action-predicting neurons under saline and DCZ treatment.

During saline control sessions, consistent with previous studies, we found that a portion of neurons encode inhibitory control, as evidenced by a significant firing rate difference between successful and failed trials (**Figure 6e-h; Supplemental figure 10**) ^4^; we therefore termed these neurons encoding neurons as their firing rates encode inhibitory control. The encoding neurons were evenly distributed across all layers of the orbitofrontal cortex (**Figure 6h**, inset). However, inhibition of LC neurons that project to the basal forebrain reduced the number of encoding neurons (**Figure 6e**; p <0.05, paired t-test). This decrease was not due to a potential reduction in the total number of neurons during DCZ sessions, as there was no significant change in the number of neurons recorded between saline and DCZ treatment sessions (**Figure 6f**; p = 0.60, paired t-test). While inhibition of BF-projecting LC neurons reduced the firing rate of these encoding neurons, it did not affect the ratio of encoding neurons with a higher firing rate in successful trials to those with a higher firing rate in failed trials (**Figure 6h**).

We also found that a small portion of neurons rapidly increased their firing rate just before action in failed trials; we therefore termed these neurons action-predicting neurons (**Figure 6i, j**). The action-predicting neurons were predominantly located in the superficial layers of the orbitofrontal cortex (**Figure 6j**, inset). In contrast to the encoding neurons, inhibition of LC neurons that project to the basal forebrain did not change the firing rate of the action-predicting neurons (p=0.40, t-test), nor did it affect the percentage of action-predicting neurons (**Figure 6k**, p=0.41, paired t-test). Moreover, a very small number of neurons were both action-predicting and encoding neurons; however, this overlap was not affected by inhibition of LC neurons that project to the basal forebrain (**Figure 6l**; p=0.43, paired t-test). Together, these results suggest that action-predicting neurons are unlikely to contribute to cognitive control of impulsivity.

We applied a stringent threshold to identify encoding neurons (p < 0.05/number of all neurons, per Bonferroni multiple comparison correction; see **Methods**), resulting in approximately 26% of neurons being classified as encoding neurons. However, it is possible that other neurons also collectively, though weakly, contributed to inhibitory control. To investigate how prefrontal population activity represents inhibitory control, we first calculated and compared pairwise cross-correlation among encoding neurons between successful and failed trials, both with and without inhibition of LC neurons that project to the basal forebrain. The inhibition of LC neurons that project to the basal forebrain more significantly disrupted the correlation structure across the encoding neurons than among the non-encoding neurons, suggesting that NE-ACh phase synchrony exerts a stronger influence on encoding neurons compared to non-encoding neurons (**Figure 7a, b**).

**Figure 7.**
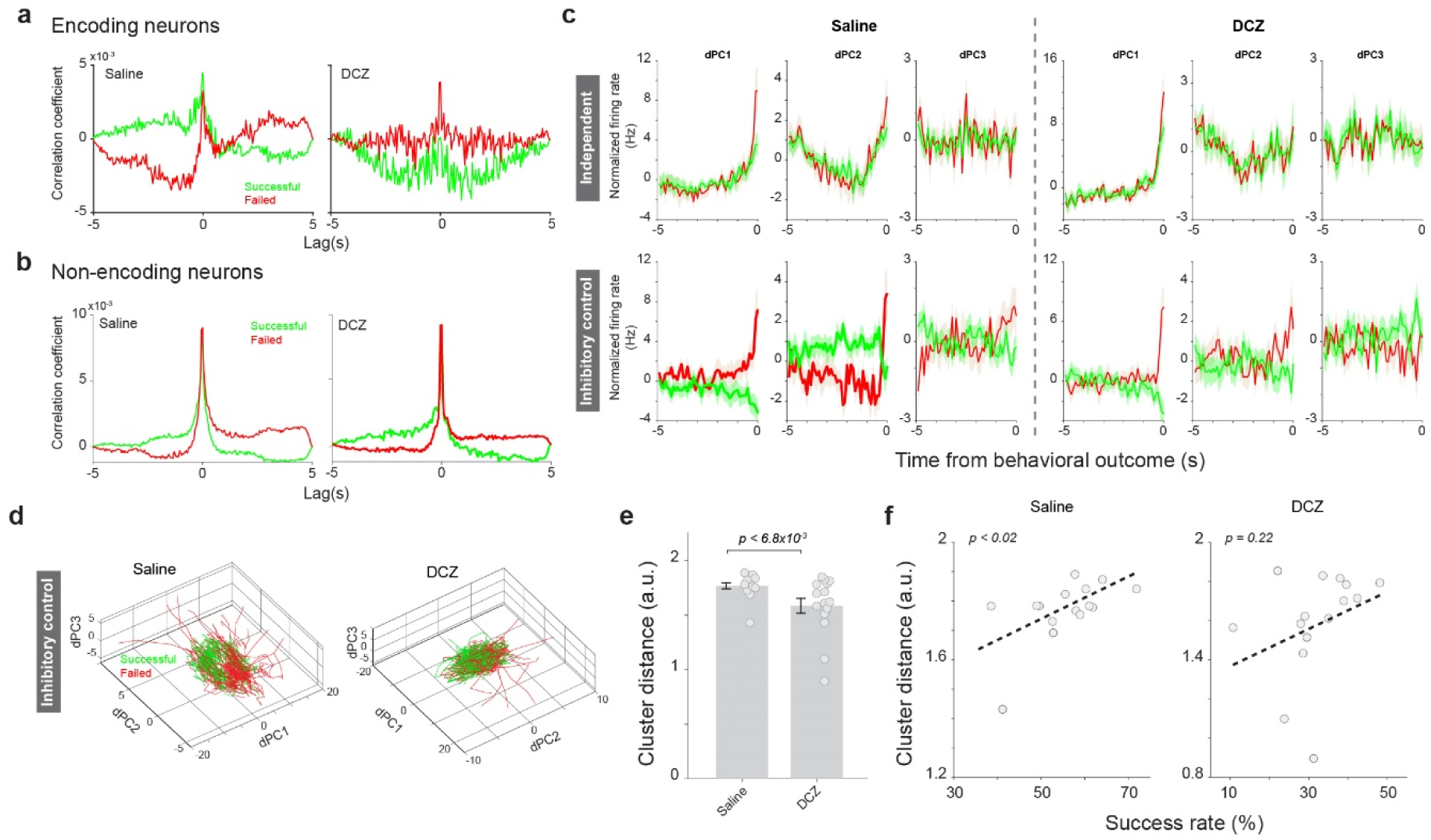
Population firing patterns encoding inhibitory control in the prefrontal cortex. **a)** Pairwise cross-correlogram across encoding neurons. 15 saline sessions and 15 DCZ sessions from 3 animals for all neuropixels results. **b)** Pairwise cross-correlogram across non-encoding neurons. **c)** Projection of population firing patterns associated with inhibitory control and independent component onto dPCI, dPC2 and dPC3, respectively. Left: saline control; right: DCZ treatment. **d)** Population firing patterns associated with inhibitory control plotted in a low-dimensional space. Left: saline control; right: DCZ treatment. **e)** Cluster distance between population firing patterns associated with inhibitory control in successful and failed trials under saline and DCZ treatment. **f)** Cluster distance between population firing patterns associated with inhibitory control in successful and failed trials is positively correlated with behavioral performance in saline control sessions (left), but not in DCZ treatment sessions (right).

To further confirm this finding, we employed demixed PCA (dPCA) analysis to decompose population dynamics associated with inhibitory control and the independent component ^44^. There is a marked difference in their projections to dPC1, dPC2, and dPC3 between successful and failed trials for population firing patterns associated with inhibitory control, but not for those associated with the independent component (**Figure 7c**). We then calculated the distance in the low-dimensional dPCA space between population firing dynamics in successful and failed trials, and found that this distance is proportional to behavioral performance (**Figure 7d-f**). To test whether non-encoding neurons contribute to the population firing dynamics representing inhibitory control, we repeated dPCA analysis using only encoding neurons or only non-encoding neurons. The distance of population firing dynamics representing inhibitory control between successful and failed trials, calculated using only encoding neurons, was significantly smaller than that calculated using all neurons (**Supplemental Figure 11**, p<8.6e-4, paired t-test), suggesting that non-encoding neurons may also contribute to inhibitory control. As expected, inhibiting LC neurons that project to the BF significantly decreased the distance between population firing patterns in successful and failed trials (**Figure 7e**, p < 6.8e-3, t-test), and further disrupted their correlation with behavioral performance (**Figure 7f**). Together, these results suggested that NE-ACh interplay in the prefrontal cortex influences the population neuronal activity that mediates inhibitory control.

#### Pupil dynamics index NE-ACh phase synchrony in failed, but not successful trials

Since previous work has demonstrated the causal relationship between LC activity and pupil dilation, as well as the correlation between pupil size and cortical NE/ACh activity ^32,33,35^, we explored the quantitative relationship between pupil size and NE/ACh as well as their phase synchrony in inhibitory control. When looking at the pupil dynamics during the inhibition tone period right before behavioral outcomes, we found an inconspicuous difference between successful and failed trials, except for the fact that the pupil started to dilate ∼0.5s before the licking in failed trials (**Figure 8a**, p=0.8, paired t-test). Given the significant difference in phase synchrony between successful and failed trials, we hypothesized that the relationship between pupil size and phase synchrony may depend on behavioral outcomes. To test this, we employed system identification approaches to assess the temporal response function (TRF, i.e. kernel) that translates NE, ACh, or phase synchrony to pupil size ^45^ (**Figure 8b**). As expected, when examining the relationship between pupil size and NE/ACh signals during the inhibition control period, we failed to find significant differences in the TRFs between successful and failed trials (**Figure 8c**). Interestingly, while the TRF mapping phase synchrony to pupil size during the inhibition control period in failed trials was similar to that of NE and ACh, the TRF in successful trials was around 0, indicating that the phase synchrony had much stronger effects on pupil size in failed trials than successful trials (**Figure 8d**). To further confirm that the difference was related to inhibitory control, we estimated the TRFs during the free period. As expected, the difference in TRFs mapping phase synchrony to pupil size between successful and failed trials was not significant (**Supplemental figure 12a**, p=0.35, paired t-test). Similarly, the TRFs that map NE and ACh signals to pupil signals in successful and failed trials were also not significantly different from each other (**Supplemental figure 12b,** NE: p=0.9, paired t-test; ACh: p=0.67, paired t-test). Additionally, their amplitude was much less than during the inhibitory control period (**Supplemental figure 12b**).

**Figure 8.**
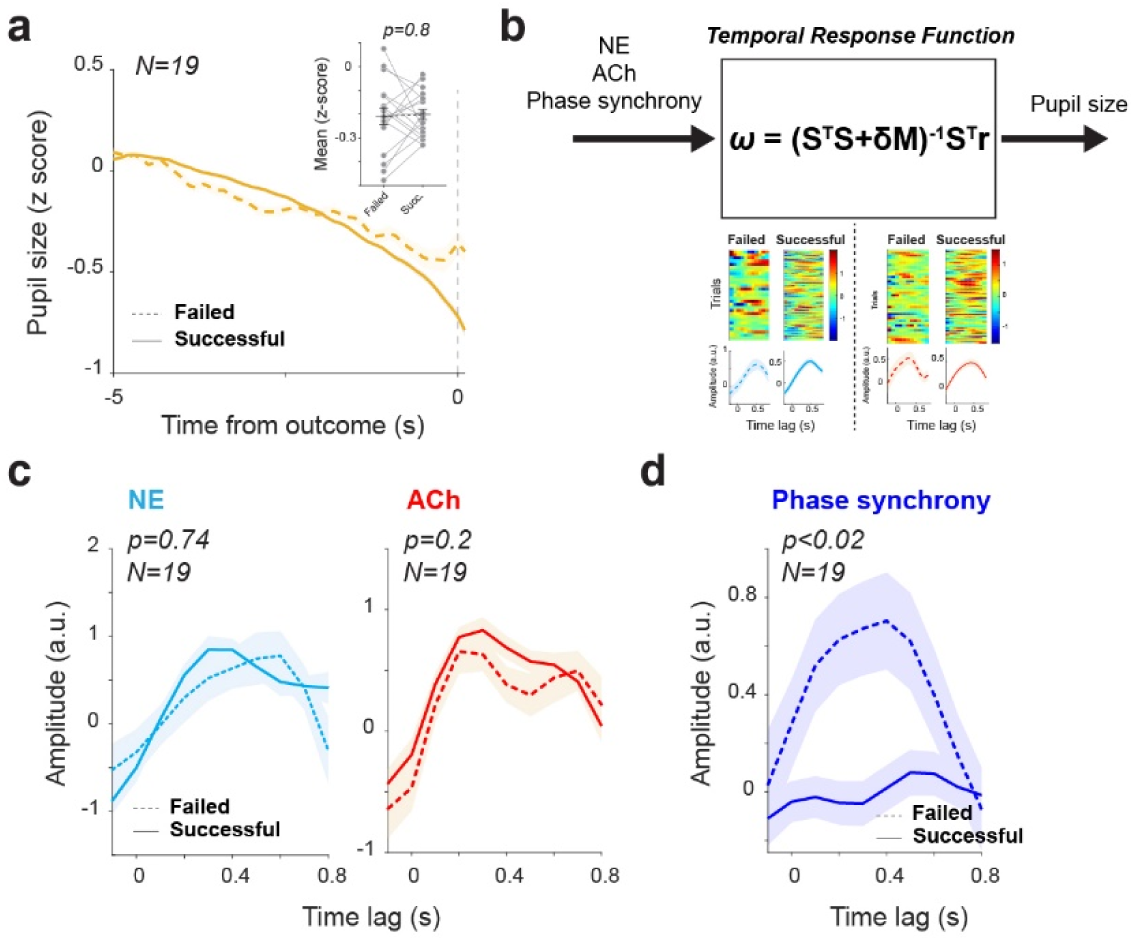
The relationship between NE/ACh phase synchrony and pupil size. **a)** Pupil dynamics prior to behavioral outcomes in the successful and failed trials of the inhibitory control task. 111 sessions from 19 animals. **b)** Identification of temporal response functions that map NE, ACh, or NE-ACh phase synchrony to pupil size. **c)** There was no significant difference in temporal response functions that map prefrontal NE and ACh signals to pupil size between the successful and failed trials in the inhibitory control task. 111 sessions from 19 animals for panels d-e. **d)** Temporal response functions mapping prefrontal NE-ACh phase synchrony were more pronounced in the failed trials than in successful trials in the inhibitory control task.

## Discussion

In the present study, we investigated the effect of synergistic activity of the noradrenergic and cholinergic systems on impulsivity control. While it has long been stipulated that the NE and ACh systems synergistically modulate brain functions, how the two systems interact to influence neural functions and behavior remains elusive. Through fiber photometry recording of NE and ACh dynamics and Neuropixels recording of population neuronal activity in the prefrontal cortex, our study, for the first time, provides direct experimental evidence supporting that the phase synchrony between the NE and ACh signals is an important neuromodulatory feature and indexes the collective effect of the noradrenergic and cholinergic systems on prefrontal neural activity mediating inhibitory control of impulsive actions.

Our findings suggested that the impulsive licking behavior observed in our mice was habitual, possibly incentivized by sweet water rewards. Moreover, the suppression of impulsive licking is a learned behavior that involves cognitive control, as evidenced by the increased success rate across training sessions (**Supplemental figure 2a**). This notion falls within the concept of inhibitory learning, which involves learning the conditions under which a response does not lead to a desired outcome and should therefore be withheld, such as not walking at a crosswalk when the light is red ^46^. The phenomenon of impulsive licking of a water spout in water-deprived rodents has been well documented ^47^, and it has been commonly used to study the neural basis of impulsivity ^47,48^. The trait of impulsivity has a multidimensional nature. It is manifested in several behavioral forms involving actions inappropriate to the situation and/or prematurely executed that often result in undesired consequences. One form of impulsivity is reflected in deficit in motor inhibition (impulsive action). This can manifest as an inability to withhold a motor response for a specified period, known as waiting impulsivity, or an inability to cancel an ongoing motor response, referred to as stopping impulsivity ^1,2^. In contrast, the other forms of impulsivity are related to impaired decision making, either resulting from inadequate evidence accumulation or due to tendency to accept immediately available but small rewards over larger but delayed rewards (impulsive choice)^2,49^. Different behavioral tasks have been implemented to assess these distinct behavioral forms of impulsivity. For instance, the probability discounting task and the temporal discounting task are mainly used to examine impulsive choice. In contrast, the stop signal reaction time (SSRT) task and the 5-choice serial reaction time (5CSRTT) task are used to examine impulsive actions^50,51^. Our behavioral task was designed to explicitly assess cognitive control of waiting impulsivity, whereas the 5CSRTT task measures both waiting impulsivity and general attentional abilities of the subjects^50^. Our study focused on prefrontal inhibitory control of waiting impulsivity and shares similarities with the delayed-response task that was used by Narayanan and Laubach ^52^, in which animals were required to hold a lever and wait until the presence of a sensory signal to release the lever. Our loss-of-function results from inactivation of the prefrontal cortex confirmed their findings that the inactivation of the rat prefrontal cortex with muscimol significantly increased impulsivity, underscoring the crucial role of the prefrontal cortex in cognitive control of impulsive behavior.

We found that a portion of prefrontal neurons encode behavioral outcomes of inhibitory control as their firing rate differed significantly between successful and failed trials. This is consistent with previous findings demonstrating distinct activation patterns among projection-specific mPFC neurons during inhibitory control ^4,47^. However, we found that these encoding neurons were evenly distributed in the mPFC and OFC, suggesting that both regions contribute to behavioral inhibition. This is consistent with previous work showing that the same behavior-relevant variable was encoded by neural activity in both mPFC and OFC ^53^. It was initially puzzling that the inhibition of LC neurons projecting to BF altered the number of encoding neurons but not the difference in their firing rate between successful and failed trials (**Supplemental Figure 10**). However, our further analysis incorporating the firing patterns of all neurons into dPCA analysis revealed that the difference in inhibitory control-related population dynamics between successful and failed trials was indeed correlated with behavioral performance. Further, inhibition of LC neurons that project to the basal forebrain diminished the difference in population firing patterns associated with two behavioral outcomes in the low-dimensional space (**Figure 7e**), suggesting a population code for inhibitory control in the prefrontal cortex. Our findings invite future investigations on how population neurons in the different regions of the prefrontal cortex mediate inhibitory control.

Previous studies utilizing pharmacological manipulations have underscored the behavioral significance of the synergy between the noradrenergic and cholinergic systems^54,55^. During the postnatal critical period, ocular dominance development in the cortex remained unaffected by the pharmacological depletion of either NE or ACh; however, combined depletion of both NE and ACh impeded this process^54^. The antidepressant effects of guanfacine, an α2 adrenergic receptor agonist, were blocked by the knockdown of nicotinic cholinergic receptor β2 subunits in the amygdala ^55^. The NE and ACh interaction in the amygdala appeared to be reciprocal because ablation of NE terminals in the amygdala also blocked the antidepressant effects of cytisine, a nicotinic partial agonist. By simultaneously measuring NE and ACh dynamics in the brain, our results offered new supporting evidence for the importance of NE-ACh interactions. Moreover, the newly developed biosensors allowed us to characterize the interaction at a behavior-relevant time scale. Our data suggested that the effect of the NE-ACh interaction on behavior is rapid because the NE-ACh phase synchrony began to decrease approximately 3 seconds before behavioral outcomes in failed trials.

Our results indicated that the NE-ACh phase synchrony, rather than individual NE or ACh signals, serves as an important biomarker for inhibitory control, highlighting the synergistic effect of NE and ACh on brain functions. Why does the NE-ACh phase synchrony matter? This may be because the interaction between the noradrenergic and cholinergic systems also happens at the receptor level. Previous work has reported that NE suppressed the release of ACh from cholinergic axonal terminals ^56^ and ACh modulation of cortical and hippocampal neurons was dependent upon NE levels^57,58^. Reciprocally, ACh inhibits NE release through M2 muscarinic cholinergic receptors^59^. It has then been argued that the balance between the noradrenergic and cholinergic systems is essential for cognitive functions^60^. In support of this notion, it has been shown that behavioral impairments in a memory task resulting from blockade of ACh inputs to the hippocampus were alleviated by a reduction in NE in the hippocampus ^60^. Therefore, it is likely that the brain operates in the optimal state when NE and ACh systems are in a certain phase relationship. This may be especially relevant given that adrenergic receptors (ARs) and muscarinic cholinergic receptors (mAChRs) are G-protein coupled receptors (GPCRs), which have slow time constants ranging from hundreds of milliseconds to seconds. In addition, among all subtypes of ARs and mAChRs, α2A AR and M1 mAChRs are the most abundantly expressed in the PFC. The inhibitory nature of α2A ARs and excitatory nature of M1 mAChRs probably make the PFC network sensitive to the phase synchrony between slow NE and ACh oscillations (0.4 – 0.8 Hz) to maintain an excitation and inhibition (E-I) balance for optimal cognitive processing ^61^. Infra-slow oscillations of LC activity have been implicated in sleep and other brain functions ^62,63^. Our data also indicate that NE-ACh phase synchrony rapidly decreases approximately 3-4 seconds prior to impulsive licking, while phase synchrony gradually increases in successful trials (**Figure 4d**). Because our data also demonstrate that the difference in phase synchrony between successful and failed trials was correlated with inhibitory control performance (**Figure 4g**), and given the reciprocal connection between the PFC and LC, it is likely that the direction of the deviation of phase synchrony from baseline during inhibitory control reflects the stability of the LC-PFC-LC network, which is essential for cognitive control. If this is the case, prefrontal NE-ACh phase synchrony may play a critical role in many other cognitive functions. It is intriguing for future work to elucidate the mechanisms through which NE-ACh phase synchrony modulates population neuronal activity mediating various executive functions at the network, cellular, and molecular levels in awake, behaving animals.

We observed robust labeling in the LC neurons upon retrograde AAV injection into the basal forebrain region. Functionally, silencing the basal forebrain-projecting LC neurons impaired inhibitory control and disrupted behaviorally relevant phase synchrony between the NE and ACh systems. These results provided new experimental evidence supporting previous work showing LC projections to the basal forebrain region^64,65^. Moreover, we improved the specificity of the AAV-mediated retrograde labeling in the present study by taking advantage of the Cre/Lox system to limit the expression of fluorescent tags to noradrenergic neurons, further supporting the notion of interaction between the noradrenergic and cholinergic systems in cognitive tasks^13,31,55,66^. However, we failed to observe evident expression of mCherry in cholinergic neurons in the basal forebrain region following retrograde AAV injection in the LC^67^. This is somewhat unexpected considering the presence of cholinergic axons in the LC area ^68^ and many previous retrograde/anterograde tracing studies reporting axonal projections from the basal forebrain to the LC region ^39,40^. One possibility is that the overserved cholinergic axons in the LC do not originate from cholinergic neurons in the basal forebrain. In support of this concept, we observed co-localization of mCherry expression and anti-ChAT signals in the trigeminal motor nucleus (V), the pontine reticular nucleus (PRN), and the superior cerebellar peduncle (SCP)/Red nucleus (RN) area^69^. Previous work has shown projections from PRN to the LC ^43^. Although the functional consequence of this cholinergic modulation of LC activity remains unclear, it could account for the slowdown of reaction time when DREADD-mediated inhibitions silenced neurons in these motor function related nuclei. Although the inhibition of these cholinergic neurons did not appear to disrupt inhibitory control functions, future work is warranted to explore the anatomy and function of these microcircuits.

Although we did not find direct projections from cholinergic neurons in the BF to the LC, it does not necessarily mean an absence of BF influence on LC activity. Cholinergic neurons in the basal forebrain could indirectly influence LC. In addition to cholinergic neurons, the basal forebrain contains two other distinct types of neurons, i.e. glutamatergic, and GABAergic^70^. It has been reported that basal forebrain cholinergic neurons that project to the prefrontal cortex have extensive local collaterals arborizing on other non-cholinergic neurons within the basal forebrain, suggesting that these neurons make local synaptic connections^70,71^. Agostinelli et al. ^72^ performed Cre-dependent anterograde tracing to investigate the targets of axonal projection of cholinergic, glutamatergic, and GABAergic neurons of the basal forebrain. Interestingly, in agreement with our data, they also failed to observe projections to the LC from cholinergic neurons in the BF area. However, they identified light projections from BF glutamatergic neurons to the LC while BF GABAergic neurons sent out moderate projection to the LC. Therefore, BF cholinergic neurons projecting to the prefrontal cortex may influence the LC through disynaptic connections to the LC. Moreover, the prefrontal cortex sends heavy projection to the GABAergic neurons, but not cholinergic neurons, in the basal forebrain ^73^. The reduced distance in prefrontal population firing patterns encoding inhibitory control between successful and failed trials, resulting from the inhibition of LC neurons projecting to the basal forebrain, may contribute to the increased difference in ACh levels between successful and failed trials through this disinhibition circuitry. We did not observe any effect of inhibiting LC neurons projecting to the basal forebrain on the differences in prefrontal NE levels between successful and failed trials. This may be because prefrontal neurons project to both the noradrenergic neurons of the LC and the GABAergic neurons surrounding it ^23^. Future work using anterograde polysynaptic viral tracers and simultaneous multi-site recordings could provide valuable insights into the neural circuitry through which the prefrontal cortex interacts with the LC or basal forebrain.

Consistent with previous findings showing the causal relationship between LC activation and pupil dilation^33,35^, we found a positive correlation between pupil size fluctuations and NE dynamics during inhibitory control. We also observed a positive correlation between pupil size and ACh signals ^32^. This could be because LC activity not only activates the BF but also dilates pupil, or due to a possible causal relationship between BF activation and pupil size. Our results showed that their relationship with pupil size was about the same between successful and failed trials for NE and ACh signals. Interestingly, the NE-ACh synchrony has a positive relationship with pupil size in failed trials but has no relationship with pupil size in successful trials. This suggested that the relationship between the NE-ACh phase synchrony is behavioral context dependent and may be gated by other cognitive-related signals. Previous work has established the causal relationship between phasic DRN activation and pupil dilation^74^. The serotonergic system has also been implicated in impulsivity control ^6,75^. Future investigations are needed to explore the dynamic relationship between pupil dilation and the collective activity of different neuromodulatory systems ^76–79^.

## Methods

All experimental procedures were approved by the Columbia University Institutional Animal Care and Use Committee (IACUC) and were conducted in compliance with NIH guidelines. Adult mice of both sexes (22 males and 26 females), aged 3 ∼ 7 months, were used in the experiments. All mice were kept under a 12-hour light-dark cycle.

### Surgical procedures

Animals were anesthetized with isoflurane in oxygen (5% induction, 2% maintenance) and secured in a stereotaxic frame. Body temperature was maintained at 36.6 ℃ using a feedback-controlled heating pad (FHC, Bowdoinham, ME). Once the animal’s condition stabilized and before an incision was made on the scalp, lidocaine hydrochloride and buprenorphine (0.05 mg/kg) were administered subcutaneously to ensure analgesics were on board during the whole surgery. At the conclusion of the surgery, Baytril (5 mg/kg) and Ketoprofen (5 mg/kg) were administered. Four additional doses of Baytril and two additional doses of Ketoprofen were provided every 24 hours after the surgery day. Animals’ weight was measured at least once per day for 5 days.

For all adeno-associated viral vector (AAV) injections, we first leveled the animal’s head by ensuring that the left and right z coordinates for the lateral scalp were within +/- 0.04 mm and the z coordinate of lambda was within +/- 0.06 mm of bregma. Burr holes were then made to target multiple brain regions, and saline was applied to each craniotomy to prevent exposed brain surface from drying out. Pulled capillary glass micropipettes (Drummond Scientific, Broomall, PA) were back-filled with AAV solution and injected into the target brain regions at 0.8 nL/s using a precision injection system (Nanoliter 2020, World Precision Instruments, Sarasota, FL). The micropipette was left in place for at least 10 minutes following each injection and then slowly withdrawn. To measure NE and ACh dynamics during inhibitory control, GRAB_NE_ (AAV9-hSyn- NE2h) and GRAB_ACh_ (AAV9-hsyn-Ach4.3) AAVs were injected into the prefrontal cortex (AP: +2.3 mm, ML: 1.2 mm, DV: -2 mm) of both hemispheres (240 nL each AAV, one AAV per hemisphere, randomly assigned, GRAB_NE_ was in the right hemisphere in 9 of 19 mice). To test cross-hemisphere similarity of NE (or ACh) dynamics, GRAB_NE_ (or GRAB_ACh_) was injected into both hemispheres (240 nL, 1 mouse for each GRAB AAV). For chemogenetic manipulation of PFC neurons, wild-type mice (RRID: IMSR_JAX:000664) were bilaterally injected with AAV9-hSyn- hM4D(Gi)-mCherry into the PFC (300 nL each hemisphere, AP: +2.3 mm, ML: 0.6∼1mm, DV: - 2.3∼-2 mm). For chemogenetic manipulation of noradrenergic or cholinergic inputs to PFC neurons, Dbh-Cre mice (RRID: IMSR_JAX:033951) or ChAT-Cre mice (RRID: IMSR_JAX:031661) were bilaterally injected with AAVrg-hSyn-DIO-hM4D(Gi)-mCherry into the PFC using the same coordinates. For chemogenetic manipulation of noradrenergic neurons projecting to the basal forebrain region, AAVrg-hSyn-DIO-hM4D(Gi)-mCherry was injected bilaterally into the BF region (300 uL per hemisphere, AP: -0.45∼0.6 mm, ML: 1.7∼1.9 mm, DV: - 4.4∼-4.2 mm) in Dbh-Cre mice ^80^. For chemogenetic manipulation of cholinergic neurons projecting to the LC region, the same retrograde AAV (300 nL per hemisphere) was injected bilaterally into the LC region (AP: -5.5 mm, ML: 0.85 mm, DV: -3∼-2.7 mm) in ChAT-Cre mice.

For retrograde labeling of LC neurons (Figure 5), retrograde AAV tracers tagged with two different fluorescent proteins were used: AAVrg-EF1a-double floxed-hChR2(H134R)-EYFP-WPRE-HGHpA (Addgene: 20298-AAVrg) and AAV-EF1a-double floxed-hChR2(H134R)-mCherry- WPRE-HGHpA (Addgene: 20297-AAVrg). These were injected into the ipsilateral PFC and BF of Dbh-Cre mice (450nL at 3 sites spanning each target region). In 2 mice, the EYFP tracer was injected into the PFC and the mCherry tracer into the BF; for the other 2 mice, this configuration was reversed. Injections were made in the left hemisphere for 2 mice and in the right hemisphere for the other 2 mice.

Following each of GRAB_NE_ and GRAB_Ach_ injections, an optical fiber (200 μm diameter & NA = 0.39 in 15 mice or 100 μm diameter & NA = 0.22 in 4 mice; RWD Life Science, San Diego, CA) was implanted with the tip of the fiber placed approximately 0.15 mm above the injection site. Bilateral optical fibers were inserted at an 8° angle ML into the PFC to create the space necessary for mounting the sleeve onto the fiber ferrules during the recordings. C&B Metabond (Parkell Inc., Edgewood, NY) was used to build a headcap to bond the fibers and a headbar. The ferrules as well as the headplate were cemented in place with dental acrylic (Prime Dental Manufacturing, Chicago, IL). Fiber photometry recording was performed 3 weeks following surgery to allow enough time for viral expression.

For Neuropixels implantation, prior to implantation, the Neuropixels 1.0 probes were mounted on a 3D-printed headcap ^81^ and the shank of the probe was stained with a solution of DiI to allow for post-mortem track localization. Mice were injected with dexamethasone (2 mg/kg), subcutaneously 2 hours before surgery to reduce swelling during surgery. After mice were anesthetized and head was leveled, a small craniotomy (1 mm diameter) was drilled over the PFC (AP: +2.3 mm, ML: 0.65 mm, DV: -2.0 mm, left PFC for 2 mice and right PFC for 1 mice). A burr hole for ground pin was also made over the occipital lobe and skull surface was roughed by scraping grids using the drill bits. Following the removal of dura at the craniotomy, probes were lowered to the targeting depth at 20 µm/min. After the implantation, the craniotomy was covered with bone wax, with a headcap and head bar cemented to the skull. The ground wire of the headcap was subsequently connected to the ground pin. Behavioral training was performed at least a week following surgery to allow for enough time for the animal to recover.

### Chemogenetic inactivation

Clozapine N-oxide (CNO, 1mg/kg; Hello Bio, Cat #: HB6149) or Deschloroclozapine (DCZ, 0.02 mg/kg; Hello Bio, Cat #: HB9126) dissolved in saline was injected intraperitoneally into the corresponding Cre mice to inactivate hM4D(Gi)-expressing neurons in the target region. DCZ is a more potent and metabolically stable DREADD agonist than CNO and was used in the later stage of the study ^82^. Saline of equivalent volume was administered as a control. Each behavioral session started 15 minutes after injection. For each animal, days of saline or CNO administration were randomly interleaved.

#### Histology

At the end of the study, mice were transcardially perfused with PBS followed immediately by ice-cold 4% paraformaldehyde. The brain was removed carefully and post-fixed overnight at 4 °C in 4% paraformaldehyde and then cryopreserved in a 30% sucrose (weight/volume) in PBS solution for 3 days at 4 ℃. Brains were embedded in Optimum Cutting Temperature Compound, and 30-μm coronal slices were sectioned using a cryostat. For brains injected with EYFP or mCherry retrograde tracers, 25-μm coronal slices were made from the appearance of the facial nerve to the point where the fourth ventricle reached its maximum size (covering the LC, approximately AP -5.2∼-5.9 mm from bregma). Brain slices were washed 4x in PBS and then incubated in 10% normal goat or donkey serum containing 0.5% Triton X-100 in PBS for 2 hours. This was followed by primary antibody incubation overnight at room temperature using 1:500 chicken anti-GFP primary antibody (A10262, ThermoFisher) for GFP/EYFP detection, 1:500 chicken anti-tyrosine hydroxylase (TH) (AB9702, Sigma-Aldrich) for TH detection, 1:200 rabbit anti- dopamine β hydroxylase (Dbh) (EPR20385, Abcam #ab209487) for Dbh detection and 1:300 rabbit anti-choline acetyltransferase (ChAT) (PA5-29653, ThermoFisher) for ChAT detection. On the next day, slices were washed 3x in PBS + Tween (0.0005%) solution followed by secondary antibody incubation for 2 hours at room temperature. For GFP/EYFP fluorescence amplification, we used 1:800 Alexa Fluor 488-conjugated goat anti-chicken (A11039, ThermoFisher). The same 488-conjugated secondary antibody was used to stain TH in the animals with chemogenetic manipulation of BF-projecting LC neurons (Figure 3b). For amplifying mCherry, we used1:800 Alexa Fluor 568-conjugated goat anti-rat (A11077, ThermoFisher). For staining ChAT and Dbh, we used 1:500 Alexa Fluor 647-conjugated goat anti-rabbit (A32733, ThermoFisher). The slices were then washed 3x in PBS + Tween solution and 1x with PBS only followed by cover-slipping using Fluoromount-G medium with DAPI (00-4959-52, ThermoFisher). Slices were imaged using 8X objective in a high-throughput slide scanner (Nikon AZ100) for further processing. Selected example slices were imaged using 20X under a confocal microscope (Nikon Ti2) with a spinning disk (Yokogawa CSU-W1).

For assessing co-localization of EYFP, mCherry and Dbh expression in LC neurons, each slice was imaged using Z-stack and tile scanning under the confocal microscope. Regions of interest (ROIs) encompassing the full extent of LC were manually drawn on low-magnification (10x) pre-scanned images based on Dbh signal intensity and referencing the mouse brain atlas ^83^, with ROI dimension ranging from 927 μm x 488 μm to 1804 μm x 927 μm (DV axis x ML axis). Slices collected towards the end that continuously lack detectable Dbh signal were excluded from analysis. For each slice, a composite image was generated by projecting all stacks using their maximum intensity. The contours of EYFP^+^, mCherry^+^ and Dbh^+^ somas were marked in ImageJ, and the centroid location of each soma was extracted using a custom Python script for visualization. The density of EYFP^+^, mCherry^+^ or dual-labeled somas was then calculated as the number of labeled somas divided by the total number of Dbh^+^ somas. We therefore reported the density of PFC-, BF- or dual-projecting LC neurons by averaging the density of corresponding tracers’ co-expression across animals.

#### Behavioral task

During the behavioral training, mice were water-deprived. During the inhibitory control task, sweetened water (10% sucrose in deionized water) was used as reward. The weight of each animal was tracked daily, and water supplements were provided after the daily training session to maintain their weight. During non-training days, animals were given ad libitum access to plain water.

During the behavioral task, the mouse was head-fixed and sat in an acrylic tube in a custom-made apparatus ^84^. Water rewards were delivered through a stainless steel feeding needle (FNS-22-1.5-2, Kent Scientific) placed ∼3mm posterior to the tip of the nose and ∼1mm below the lower lip. A capacitance touch sensor (AT42QT, SparkFun) was connected to the water spout to detect licks. Inhibition tone (4 kHz, 65 dB) was generated with an Arduino Mega 2560 microcontroller, sent to an audio amplifier, and played via a speaker placed 25 cm from the animal. Punishment air puffs were delivered through a 16-gauge stainless-steel tube placed approximately 8 cm from the animal’s cheek and was contralateral to the pupil camera. Control of the behavioral experiment and sampling of animals’ behavioral responses were performed by custom-programmed software running on a MATLAB xPC target real-time system (Mathworks, Natick, MA). All behavioral data was sampled at 1 kHz and logged for offline analyses.

During the shaping period, mice first went through a water-association phase during which the animals learned to lick from the water spout to collect sweetened water delivered with a random interval (12 to 22 s uniform distribution). The animal advanced to the phase 2 once it licked for >75% of water deliveries.

During the phase 2, the animals had a free period (5 to 7 s uniform distribution) at the beginning of each trial, where licking did not result in any punishment. After the free period, an inhibition tone (4 kHz, 65 dB) was played for a random period (2 to 5 s, exponential distribution, λ=1.5), during which any lick would immediately terminate the inhibition tone and trigger three low intensity air puffs (10 psi, 20 ms, 200 ms inter-puff-interval) to their cheek as a punishment. In the case where the animals withheld their licks throughout the inhibition tone, a drop of sweetened water (∼6 μL) was immediately delivered after tone ended as a reward. Because rodents usually collect water rewards within 0.8 s ^85^, the trial was considered as a successful trial if animal licked to collect the reward within 1 s (an exclusion threshold of 0.75 s or 1.25 s resulted in similar results). The trials where the animals collected water rewards outside the window of opportunity of 1 s (6% trials) were excluded from further analysis. An inter-trial period of 7 to 10 s (uniform distribution) was added to the end of each trial. Animals advanced to the full inhibitory control task once they reached a >50% success rate for at least 2 consecutive days in the Phase 2 task.

The full inhibitory control task was similar to the phase 2 task, except for a longer inhibition tone (5 to 12 s exponential distribution, λ=4.5, **Figure 2a**) and more severe punishment (an air puff of 20 psi for 200 ms) for failed trials. Animals were considered as proficient when their behavioral performance had been above the chance-level success rate (see details in Behavior analysis section) for at least 3 consecutive days. Animals whose behavioral performance was not significantly greater than the chance level were excluded from further analyses (N=1). Fiber photometry was recorded in the PFC of 19 animals and Neuropixels recordings were made in 3 animals.

#### Neuropixels recording

Recordings from Neuropixels probes were acquired using an NI Neuropixels recording system controlled by SpikeGLX (Release v20230905-phase30). The NI recording system was synchronized with the behavioral apparatus via TTLs generated by the xPC real-time system.

#### Fiber photometry recordings and preprocessing

Fluorescence signals mediated by the GRAB_NE_ and GRAB_ACh_ sensors were recorded using a 2-channel fiber photometry system (Doric Lenses). For the recording of each sensor, excitation lights with 465 nm and 405 nm wave lengths were generated by LEDs (CLED_465 and CLED_405, Doric Lenses) and passed through a MiniCube (iFMC4_AE (405)E(460–490)_F(500– 550)_S, Doric Lenses). Emission fluorescence from the GRAB_NE_ and GRAB_ACh_ sensors was measured by an integrated PMT detector in the MiniCube. The fiber photometry recordings were run in a ‘Lock-in’ mode controlled by Doric Neuroscience Studio (V5.4.1.12), where the intensity of the four excitation lights was modulated at frequencies of 208.62 Hz, 572.21 Hz, 333.79 Hz, and 470.88 Hz, respectively, to avoid contamination from other light sources in the room and crosstalk between the excitation lights. The demodulated signal processed by the Doric fiber photometry console was low-pass filtered at 25 Hz and sampled at 12 kHz with a 16-bit ADC. The fiber photometry system was synchronized with the behavioral apparatus through TTLs generated by the xPC target real-time system (MathWorks, Massachusetts). All photometry data were decimated to 120 Hz by Doric Neuroscience Studio software and saved for offline analysis. Since we are interested in NE and ACh dynamics during the inhibitory control periods (average duration of the inhibition tones = 7.6 s), fluorescence signals were first high-pass filtered (cutoff frequency, 0.1 Hz) to remove the low-frequency oscillations and then used to calculate NE and ACh dynamics during the inhibitory control.

#### Pupillometry

Pupil recordings were obtained using a custom pupillometry system ^86^. The camera was triggered by 10-Hz TTLs from the xPC target real-time system. Pupil images were streamed to a high-speed solid-state drive for offline analysis. For each video clip, a region of interest (ROI) was manually selected initially. The DeepLabCut toolbox was used to segment the pupil contour ^87^. Training sets were created, consisting of 450 frames for video clips with a resolution of 800*600 pixels or 160 frames for video clips with the resolution of 1280*1080 pixels. Within each frame, 12 points around the pupil were manually labeled, and cropping parameters were adjusted to enhance training accuracy. The mobilenet_v2_0.75 deep network was trained on each frame and employed for the analysis of video clips from all sessions. Circular regression was then applied to fit the automatically labeled points, enabling the computation of pupil size based on the fitted contour. To ensure segmentation accuracy, approximately 5% of segmented images were randomly selected and inspected. Pupil size during periods of blinks was estimated through interpolation, using pupil sizes just before and after blinks. If DeepLabCut could not recognize pupil contour due to either poor video quality or animal’s eyelid covering a significant portion of pupil in >33% of the recorded video frames, the session was excluded from pupillometry analysis. Prior to further analysis, a fourth-order non-causal low-pass filter with a cutoff frequency of 3.5 Hz was applied to the pupil size data^33,35^.

### Data analysis

All data analyses were first conducted on individual sessions. Grand averages and standard errors of means were then calculated across sessions or animal subjects for analysis and visualization. For each session, the first trial was excluded due to the time required for experimenter to leave the behavioral training room.

#### Behavior

To measure the effect of inhibitory control on suppressing impulsive licking, we normalized experimentally-measured success rate with chance-level success rate. The chance-level success rate for each animal was determined via Monte-Carlo simulations based on the animal’s baseline behavior (all sessions for WT mice (C57BL6/J, RRID: IMSR_JAX:000664) and the saline sessions for transgenic mice). In these sessions, the time stamp of licks from 7 s after the behavioral outcome of the current trial to inhibition tone onset of the next trial was used to calculate the distribution of the baseline inter-lick-intervals. As such, licks in response to water rewards/punishment were excluded from the assessment of baseline inter-lick-interval distribution. For each simulated trial, the duration of the inhibition tone was drawn from an exponential distribution from 5 to 12 s with λ=4.5, the same distribution used in the full inhibitory control task. A baseline lick sequence was simulated using the estimated baseline inter-lick-interval distributions. If a lick was detected during the inhibition tone period, a failure trial was recorded. Otherwise, a successful trial was logged. Each simulated session comprised 100 trials and chance-level success rate was calculated by averaging across 10 simulated sessions for each animal.

For each animal, normalized performance for each experimental session was calculated as below: _normalized performance =_ raw sucess rate - chance level success rate _∗ 100%_ chance level success rate

Reaction times were computed as the time from water reward onset, which marked the beginning of the window of opportunity, until the first lick response for collecting the water reward. Reaction times were only computed for successful trials.

#### Frequency analysis

For each session, NE and ACh signals were z-scored and the one-sided power spectral density (PSD) was computed using a Hamming window with a segment overlap of 50%, employing the MATLAB built-in function *pspectrum*. For **Figure 1f**, spectrum was first normalized to the power at 0.1 Hz for each session and then averaged across sessions ^88^.

#### Correlation analysis

To assess the correlation between NE eand ACh signals, both signals were band-pass filtered (cutoff frequencies, 0.1 and 5Hz) and z-scored. For each session, the overall cross-correlation between NE and ACh dynamics was calculated using the MATLAB built-in function *xcorr*, with a maximum lag of 10 s. To further evaluate the relationship between NE and ACh dynamics specifically during inhibitory control period, Pearson’s correlation coefficients were calculated between the two traces extracted from a 4-s window spanning 5 to 1 s prior to the behavior outcome across trials. To calculate the cross-correlogram between pupil size and NE/ACh signals, NE and ACh signals were first low-pass filtered (cutoff frequency 5 Hz) then down-sampled at 10 Hz (i.e. pupillometry frequency). All signals, including pupil size, were z-scored, and cross-correlations for each session were computed. Confidence intervals were generated by performing 100 iterations of cross-correlation calculations between one original signal and a shuffled version of the other signal. The 0.15th and 99.85th percentiles of the resulting distributions were used to define a 99.7% confidence.

#### Coherence analysis

To examine how closely NE and ACh interact with each other over different frequency bands, we computed coherence using a custom MATLAB script adapted from the Buzsaki laboratory (chronux, https://github.com/buzsakilab/buzcode) ^89^. The NE-ACh coherence was calculated using the function *cohgramc* in a multi-taper manner (window, 11s; overlap, 5s; step, 6s; number of tapers, 8; padding, 0) for each session, and then averaged across sessions. Because the toolbox requires an alignment of recorded data, we used the first 1500 s of each recording to calculate coherence. The confidence intervals were estimated by repeatedly calculating coherence between shuffled signals for each session.

#### Phase relationship with pupil

To evaluate the phase relationship between NE/ACh signals and pupil fluctuations, we aligned the amplitude of NE or ACh signals to different binned phases (36 bins from -180° to 180°, 10° per bin) of a canonical cycle of pupil fluctuation ^32^. To achieve this, we first band-pass filtered (cutoff frequencies, 0.1 and 1Hz) the pupil signal and calculated its instantaneous phase angles using Hilbert transform for each session. Subsequently, we identified the phase of pupil signals at each time point and then averaged NE/ACh signals aligned with each bin of the pupil phase.

#### Phase synchrony between NE and Ach

To examine the phase interplay between NE and ACh over time, instantaneous phase synchrony ^38^ between the two signals was calculated. For each session, each photometry signal was first band-pass filtered (cutoff frequencies, 0.4 and 0.8 Hz; Butterworth; order 2) and z-scored. Hilbert phase angles were then extracted and wrapped into [- 180, 180] degrees. The instantaneous phase synchrony was calculated as follows ^38^:

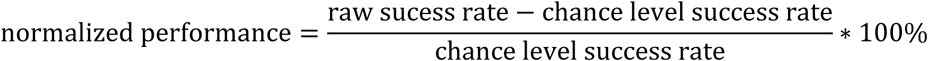

where 𝜑_NE_(𝑡) and 𝜑_ACh_(𝑡) are instantaneous phases of NE and ACh, respectively. A second-order non-causal low-pass filter with a cutoff frequency of 4 Hz was used to de-noise the calculated phase synchrony of each session prior to further analyses. Phase synchrony of 1 indicates |𝜑_NE_(𝑡) - 𝜑_ACh_(𝑡)| = 0 and thus indicates the NE and ACh oscillations are in-phase. On the contrary, phase synchrony of 0 indicates |𝜑_NE_(𝑡) - 𝜑_ACh_(𝑡)| = 180^°^ and thus indicates the NE and ACh oscillations are out-of-phase (**Figure 4a**).

To quantify the number of switching of NE-ACh relationship from in-phase to out-of-phase, we applied Hilbert transform to phase synchrony for each session and counted the number of phase-wrapping points as the number of the switching (Figure 4a). Phase synchrony prior to behavioral outcomes were averaged using a 2-s moving window with a 0.5 s step. Switching rates prior to the behavioral outcomes were estimated by first counting the number of switching event within a 2-s window then the switching rate was smoothed using a Gaussian window with a SD of 0.25.

#### Temporal response function (TRF) analysis

We applied TRFs estimation (mTRF-Toolbox v2.3, MATLAB R2018a) ^45^ to characterize the transform relationship from NE/ACh/NE-ACh phase synchrony to pupil size. Here, pupil size was modeled in a forward direction by convolving the TRF kernel with other signals. The model equation reads as follows:

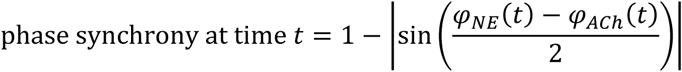

where {𝑝}_t_ represents the pupil size, {𝑠}_t_ represents NE, ACh or NE-ACh phase synchrony, and {𝛽}_r_ represents the TRF; {𝜀}_t_ is the residual response accounting for the noise not explained by the model. Therefore, the weights in 𝛽(𝜏) characterize the transformation of NE, ACh, or NE-ACh phase synchrony to pupil size for a range of time lags 𝜏.

The weights of TRF were estimated by minimizing the mean-squared error (MSE) between the measured pupil traces and those reconstructed by the convolution for each trial. Tikhonov regularization was applied to avoid overfitting ^90^

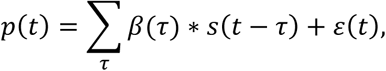

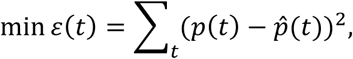

where 𝛃 and 𝐏 are vectors of TRF weights and pupil size traces, 𝐒 is the design matrix containing the time lagged signals related to neurotransmitters, 𝐈 is the identity matrix and 𝜆 is the regularization parameter. Leave-one-out-cross-validation was conducted on the traces for each trial to identify the optimal 𝜆. MSEs resulted from different 𝜆 values were averaged across left-out folds and the finalized 𝛃 of the current trial was obtained from parameters corresponding to the minimum MSE.

For estimating TRFs during inhibitory control, periods from inhibition tone onset to outcome of each trial were chosen. For estimating TRFs during lick-free period, periods from 5s prior to trial onset to inhibition tone onset of each trial were used. TRF of each session was obtained by averaging weights across trials from each of the two behavioral outcome types.

#### Discriminability analysis

We applied both the signal detection theory and machine learning algorithm to examine the discriminability between physiological signals associated with failed and successful trials ^37,91,92^. For each session, we first conducted a receiver operating characteristics (ROC) analysis using NE or ACh signals averaged over a 2-s window from 3s to 1s prior to the behavioral outcome across all trials. At each time point, the signal distributions associated with failed or successful trials was constructed and an ROC curve was created based on the distributions. In our case, the ROC curve expresses the two probabilities as follows:

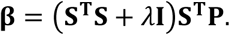

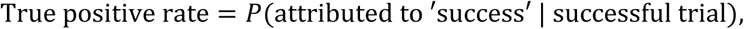

To evaluate the overall discriminability, the area under the ROC curve (AUROC) was calculated for each time point and averaged across sessions (240 time points in total). Additionally, we employed a support vector machine (SVM, *sklearn.svm* in Python) to assess the discriminability of physiological signals related to two behavioral outcomes. The NE or ACh trace within the same 2-s period were used directly as input, with the behavioral outcome being the label for each trial. For each session, an SVM classifier was trained and tested using 10-fold cross-validation. The classification performance was quantified using balanced accuracy to account for potential class imbalances ^93^:

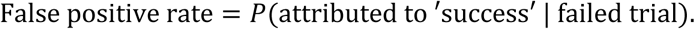

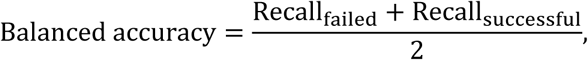

#### Neuropixels data analysis

Spikes from electrophysiological data of each recording session were automatically detected using KiloSort 3. Units labeled as ‘noise’ by the automatic spike-sorting were discarded. The spike data were then manually curated using the open-source package Phy (https://github.com/kwikteam/phy). Only units labeled as ‘good’ by the automatic spike-sorting and manual curation were included in further analysis. For the identification of regular spiking units (RSUs) and fast spiking units (FSUs), we chose the width (full-width half maximum of the post-hyperpolarization peak) and the trough to peak duration ^94^ of action potentials (AP) as our waveform features, and performed a Gaussian mixture model (GMM) hard clustering in the two-dimensional feature plane.

Because we were interested in neurons that may contribute to the cognitive control of impulsivity, we analyzed only those neurons that were active (firing rate > 1 Hz) during the inhibitory control task. Encoding cells were defined as neurons that exhibited a significant difference in firing rate between successful and failed trials while the remaining cells are non-encoding. The significance of this difference was evaluated using a paired statistical test, with a p-value set to 0.05 divided by the number of total cells in the session (i.e., Bonferroni multiple comparison correction). Action-predicting cells were defined as neurons whose firing rate was greater than 2 standard deviations above the mean firing rate within 0.3 seconds before the behavioral outcome (i.e., licking) in failed trials.

As we are interested in the correlation structure across prefrontal neurons during inhibitory control, we used the spikes recorded within 5 seconds before behavioral outcomes to calculate the cross-correlogram. The pairwise cross-correlogram was calculated by averaging the cross-correlogram across all cell pairs. We then subtracted a shuffled cross-correlogram, computed using spikes from cell pairs in randomly shuffled trials ^95^ from this raw cross-correlogram. This shuffling process was repeated 100 times to estimate the final shuffle-corrected cross-correlogram.

#### Demixed PCA (dPCA) analysis

To further characterize prefrontal neuronal population activity structure over the period prior to behavioral outcomes, we employed demixed PCA analysis. We chose dPCA over traditional PCA analysis as it allows the decomposed components to capture the maximum variance in neural population activity explained by task-related variables ^44^. In our study, dPCA decomposes the raw population neural activities into the following marginalization:

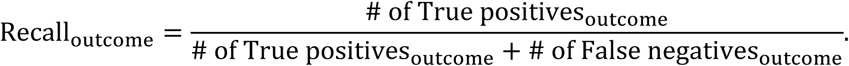

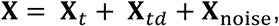

where 𝐗 is an NxDxTxK matrix containing firing rates of individual neurons within 1 to 5 seconds before the behavioral outcome in individual trials calculated using 100-ms windows. N is the number of active neurons, D is the number of inhibitory control outcomes, T is the number of time points, and K is the number of trials (for an outcome with a fewer number of trials, *NaN*s were assigned to the empty entries). 𝐗_t_ corresponds to the time effect (the Independent component in **Figure 6c**) with NxT entries replicated DxK times. 𝐗_td_ corresponds to the cognitive control effect (the Inhibitory control component in **Figure 6c**) with NxTxD entries replicated K times. 𝐗_noise_represents the noise. The loss function is given by:

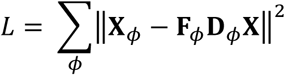

where 𝐗_<P_ is a specific decomposition, and 𝐅_<P_ and 𝐃_<P_ are the encoder and decoder matrix of that decomposition, respectively.

dPCA analysis was performed using a custom script employing functions in the dPCA package (https://github.com/machenslab/dPCA) with default parameters. For each session, we computed in total 20 dPCs (or number of neurons if that number is less than 20, for PFC neuron subgroups, i.e. non-encoding and encoding) and retained the top 3 dPCs of both ‘Independent’ and ‘Inhibitory control’ decomposition. The population firing activities were then projected onto the respective dPC decoder axis with the resulting projections being averaged across sessions. To quantify the difference in neural population dynamics between trials of different outcomes, we calculated the Mahalanobis distance in the dPCA space between population dynamics prior to success and failure ^96^. For visualizing session trajectory of population dynamics in the dPCA space, the projected trace along each dPC axis was smoothed using a 500-ms moving window. This smoothing was for visualization only and not for quantitative analysis.

#### Statistics

All statistical tests were two-sided. A one-sample Kolmogorov–Smirnov test was used to assess the normality of data before performing statistical tests. If the samples were normally distributed, a paired or unpaired t-test was used. Otherwise, the two-sided Mann–Whitney U-test was used for unpaired samples or the two-sided Wilcoxon signed-rank test for paired samples. Bonferroni correction was used for multiple comparisons. No statistical methods were used to predetermine sample sizes, but our sample sizes are similar to those in previous reports and are typical for the field. Randomization of inhibition tone duration and simulation was generated by using MATLAB random number generators. Data collection and analysis were not performed blind to the conditions of the experiments.

## Authors contribution

Q.W. designed and supervised the study. Y. Liu performed the experiments and analyzed the data. Y. Liu, P.S. and Q. W. interpreted the results. J.F., G.L. and Y. Li provided GRAB_NE_ and GRAB_ACh_ sensors. Y. Liu, P.S. and Q.W. wrote the manuscript. All authors commented on and approved the manuscript.

## Acknowledgements

The authors would like to thank M. Shadlen and J. Gottlieb for helpful discussions, C. Slater and M. Sorrentino for their assistance in constructing Neuropixels implants and experiments, and C. Martinez for assistance in confocal imaging, which was performed with support from the Zuckerman Institute’s Cellular Imaging platform.

## Disclaimer

Q.W. is the co-founder of Sharper Sense.

---

**Supplemental figure 1.**
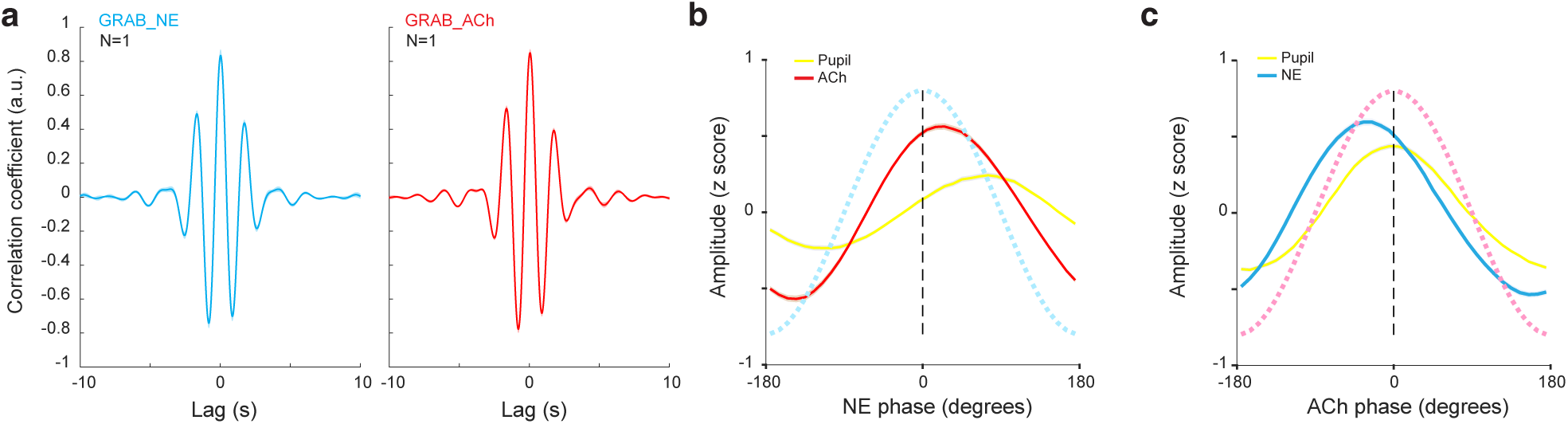
**a)** Cross-correlogram of GRAB_NE (left) and GRAB_ACh (right) signals simultaneously recorded from the PFC of both hemispheres. **b)** ACh and pupil phases referenced to the NE phase. **c)** NE and pupil phases referenced to the ACh phase.

**Supplemental figure 2.**
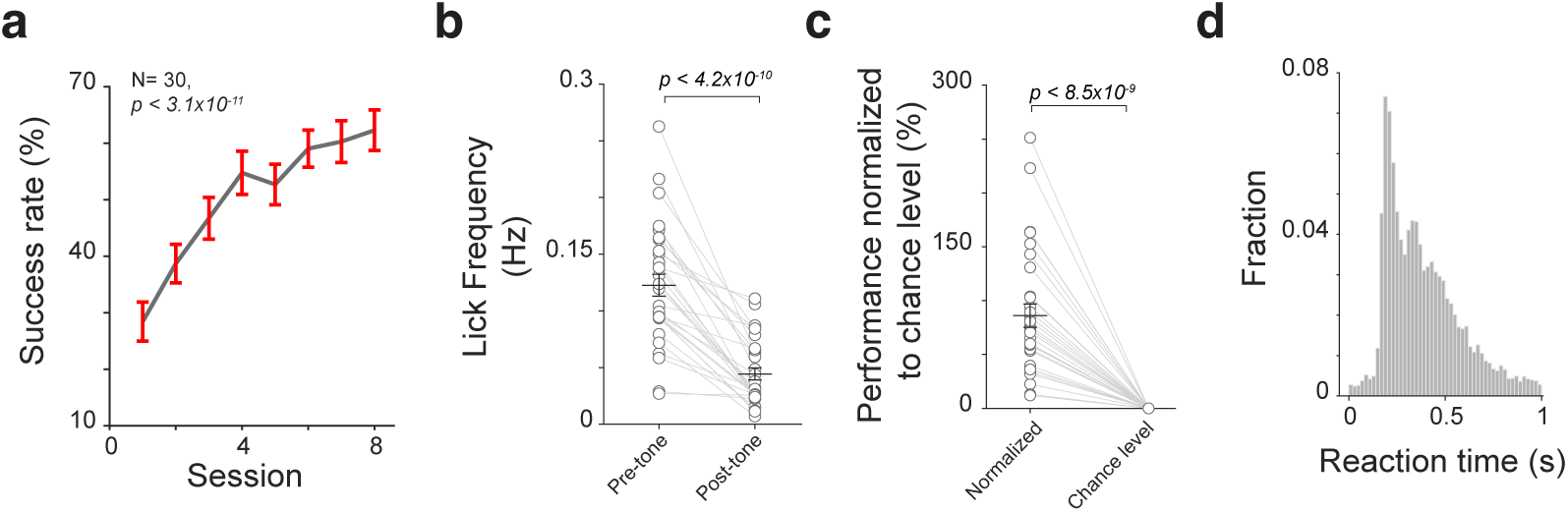
The inhibitory control task performance. **a)** The increase in success rate during the initial training sessions indicates that the suppression of impulsive licking was a learned behavior. **b)** Mean licking frequency within a 2-second window prior to vs. after the onset of the inhibition tone. **c)** Inhibitory control task performance normalized to chance-level performance. **d)** Histogram of reaction time in successful trials.

**Supplemental figure 3.**
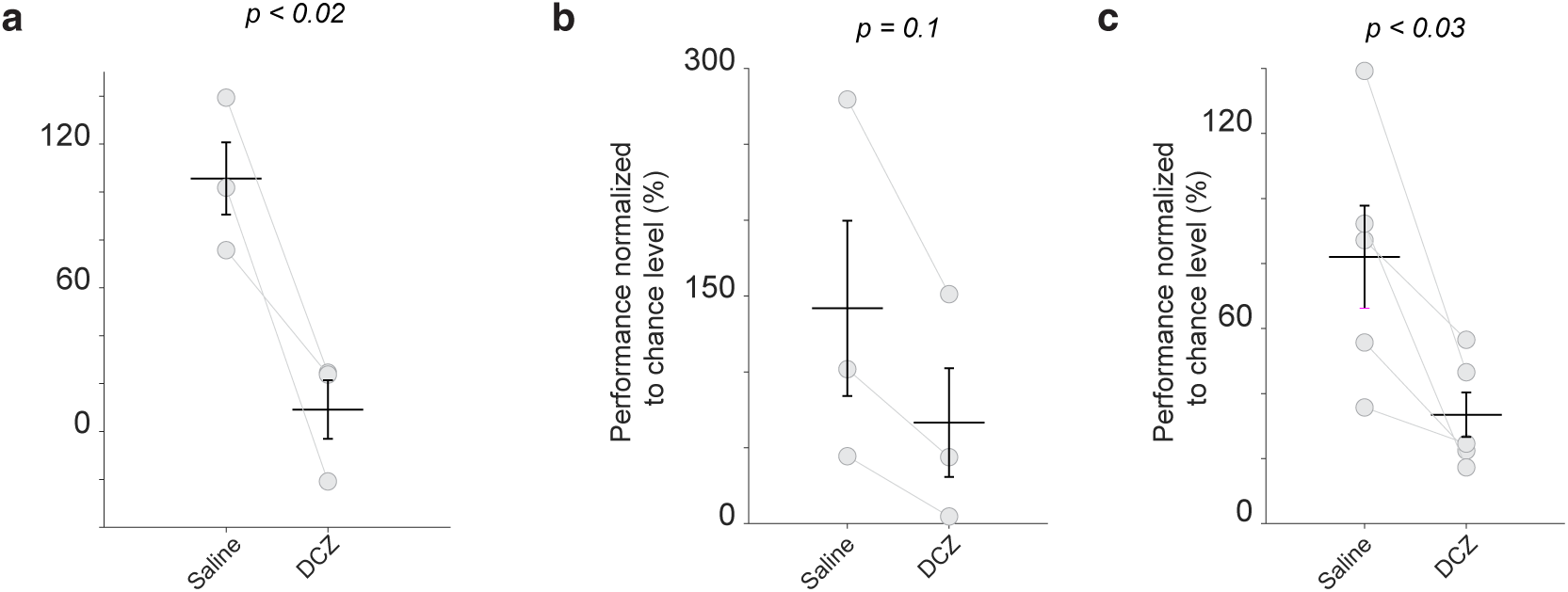
Normalized performance in loss-of-function experiments. **a)** Normalized performance with and without the inactivation of the prefrontal cortex. **b)** Normalized performance with and without the inactivation of noradrenergic inputs to the prefrontal cortex. **c)** Normalized performance with and without the inactivation of cholinergic inputs to the prefrontal cortex.

**Supplemental figure 4.**
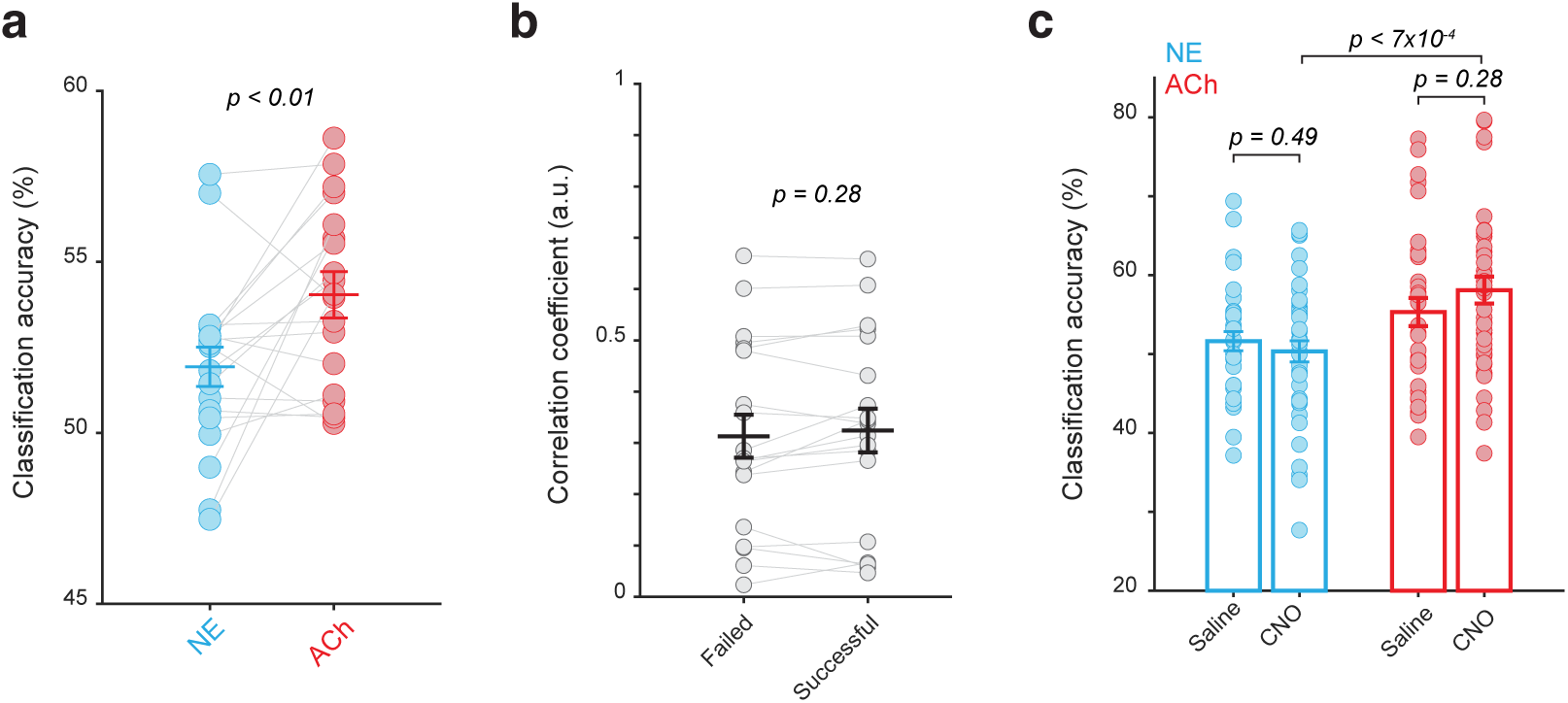
Single-trial analysis of NE and ACh dynamics during inhibitory control. **a)** Classification accuracy of a support vector machine (SVM) classifier using single-trial NE and ACh dynamics during inhibitory control. **b)** Correlation between single-trial NE and ACh dynamics in successful and failed trials. **c)** SVM classification accuracy with and without inhibition of LC neurons projecting to the basal forebrain.

**Supplemental figure 5.**
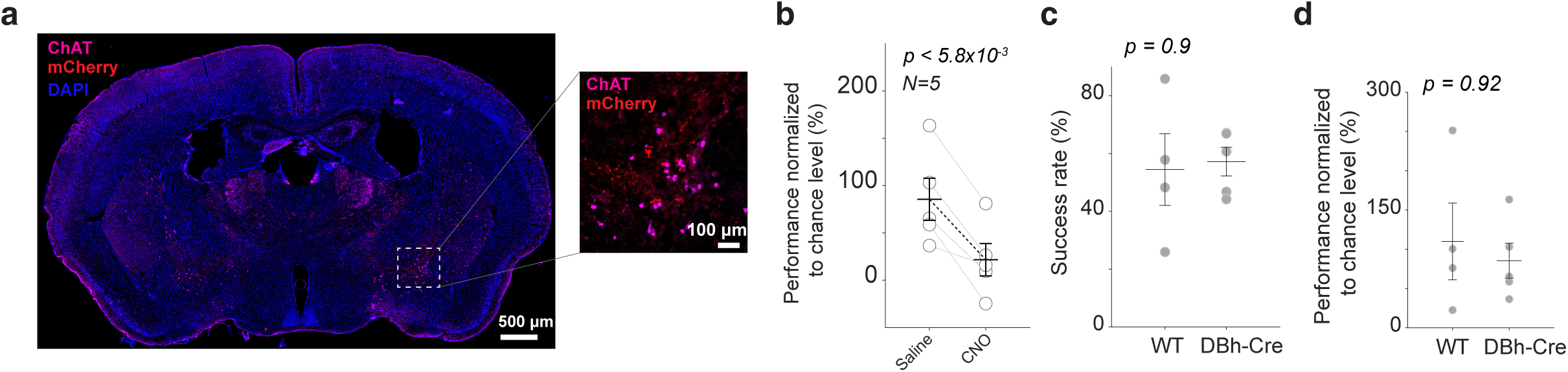
**a)** mCherry expression in putative LC axons in the basal forebrain region of DBh-Cre mice. **b)** Normalized behavioral performance with and without inhibition of LC neurons projecting to the basal forebrain. **c)** Raw success rate of WT and DBh-Cre mice in saline control sessions. **d)** Normalized behavioral performance of WT and DBh-Cre mice in saline control sessions.

**Supplemental figure 6.**
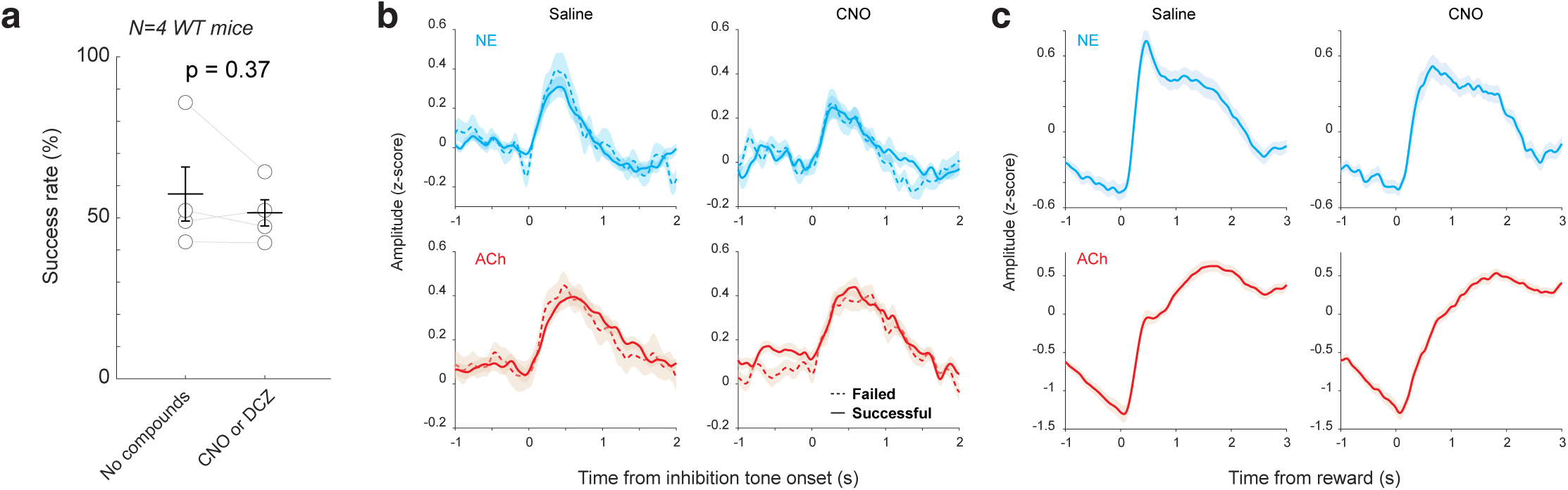
**a)** CNO alone did not affect animals’ inhibitory control. **b)** NE and ACh dynamics around the onset of the inhibition tone in successful and failed trials, with and without inhibition of LC neurons projecting to the basal forebrain. **c)** NE and ACh dynamics around water rewards with and without inhibition of LC neurons projecting to the basal forebrain.

**Supplemental figure 7.**
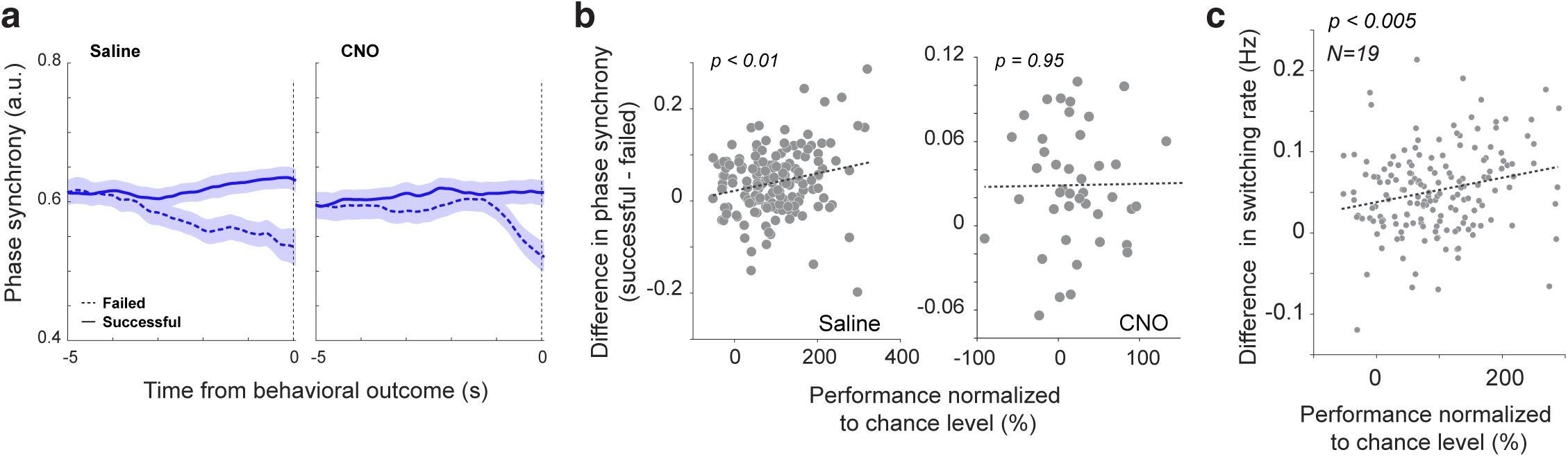
**a)** NE-ACh phase synchrony calculated using ACh signals shifted 30 ms forward. **b)** The difference in phase synchrony between successful and failed trials was positively correlated with inhibitory control performance during saline control sessions but not under CNO-mediated inhibition of LC neurons projecting to the basal forebrain. **c)** The difference in switching rate between successful and failed trials was positively correlated with inhibitory control performance.

**Supplemental figure 8.**
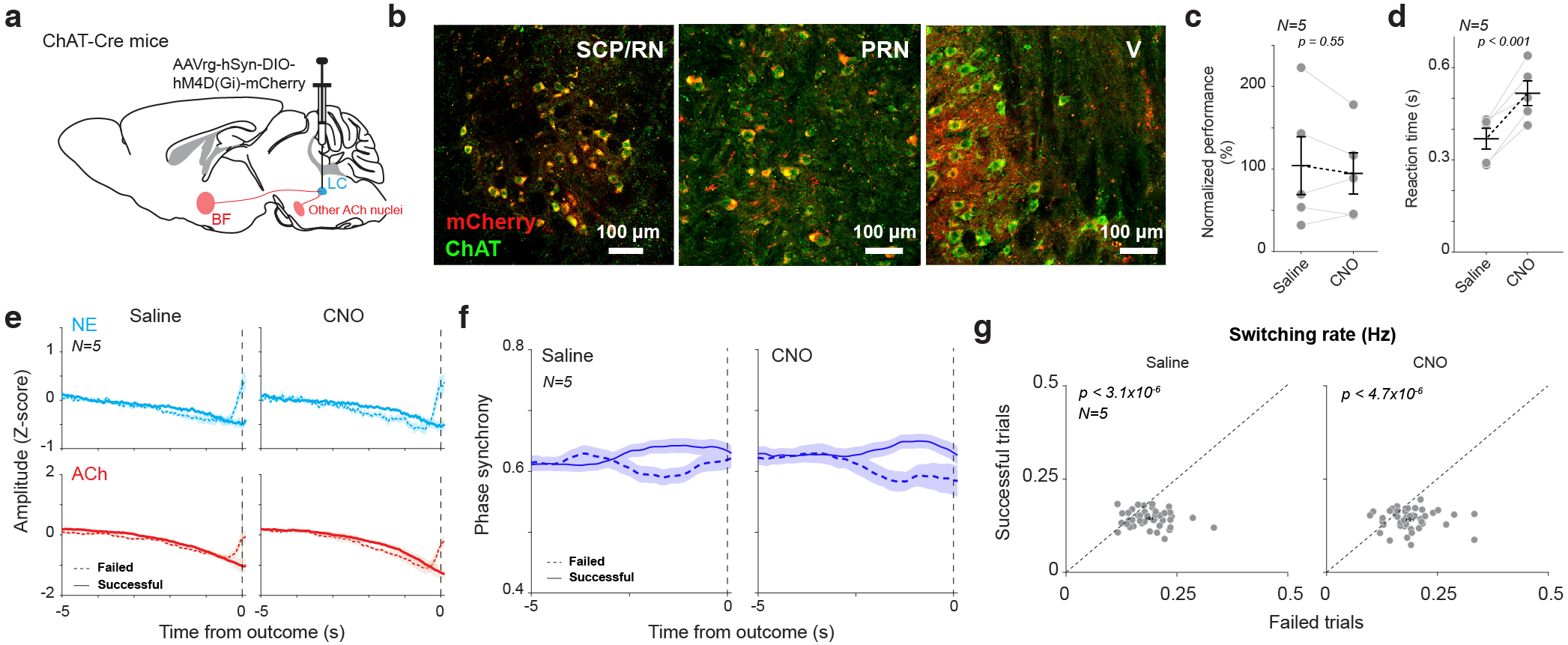
Chemogenetical/y silencing cholinergic neurons that project to the LC area did not impair inhibitory control, nor did it change prefrontal NE-ACh phase synchrony. **a)** Diagram of retrograde expression of DREADD receptors in cholinergic neurons that project to the LC area. **b)** Histological confirmation of mCherry expression in the superior cerebellar peduncle (SCP)/Red nucleus (RN), pontine reticular nucleus (PRN), and trigeminal motor nucleus (VJ. **c)** Chemogenetic inhibition of cholinergic neurons that project to the LC region had no effect on inhibitory control, **d)** Chemogenetic inhibition of cholinergic neurons that project to the LC region slowed reaction time. **e)** Prefrontal NE/ACh signals prior to behavioral outcomes in the successful and failed trials under saline or CNO treatment. **f)** Prefrontal NE-ACh phase synchrony prior to behavioral outcomes in the successful and failed trials under saline and CNO treatment. **g)** Mean switching rate prior to behavioral outcomes in the successful vs. failed trials under saline or CNO treatment. All data are from 43 saline sessions and 43 CNO sessions from 5 animals.

**Supplemental figure 9.**
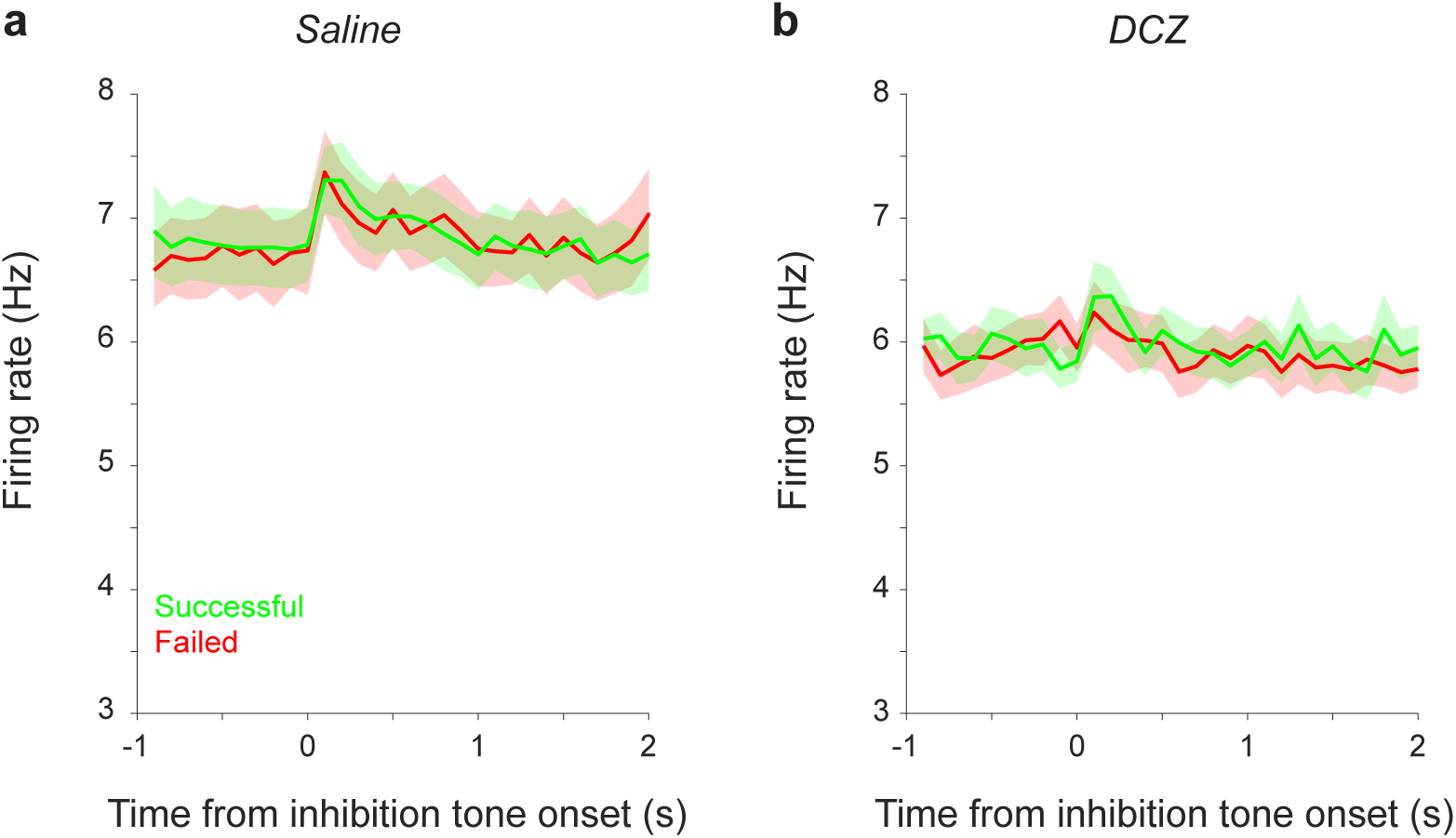
Firing rates of prefrontal neurons around the onset of the inhibition tone, with (a) and without (b) inhibition of LC neurons that project to the basal forebrain.

**Supplemental figure 10.**
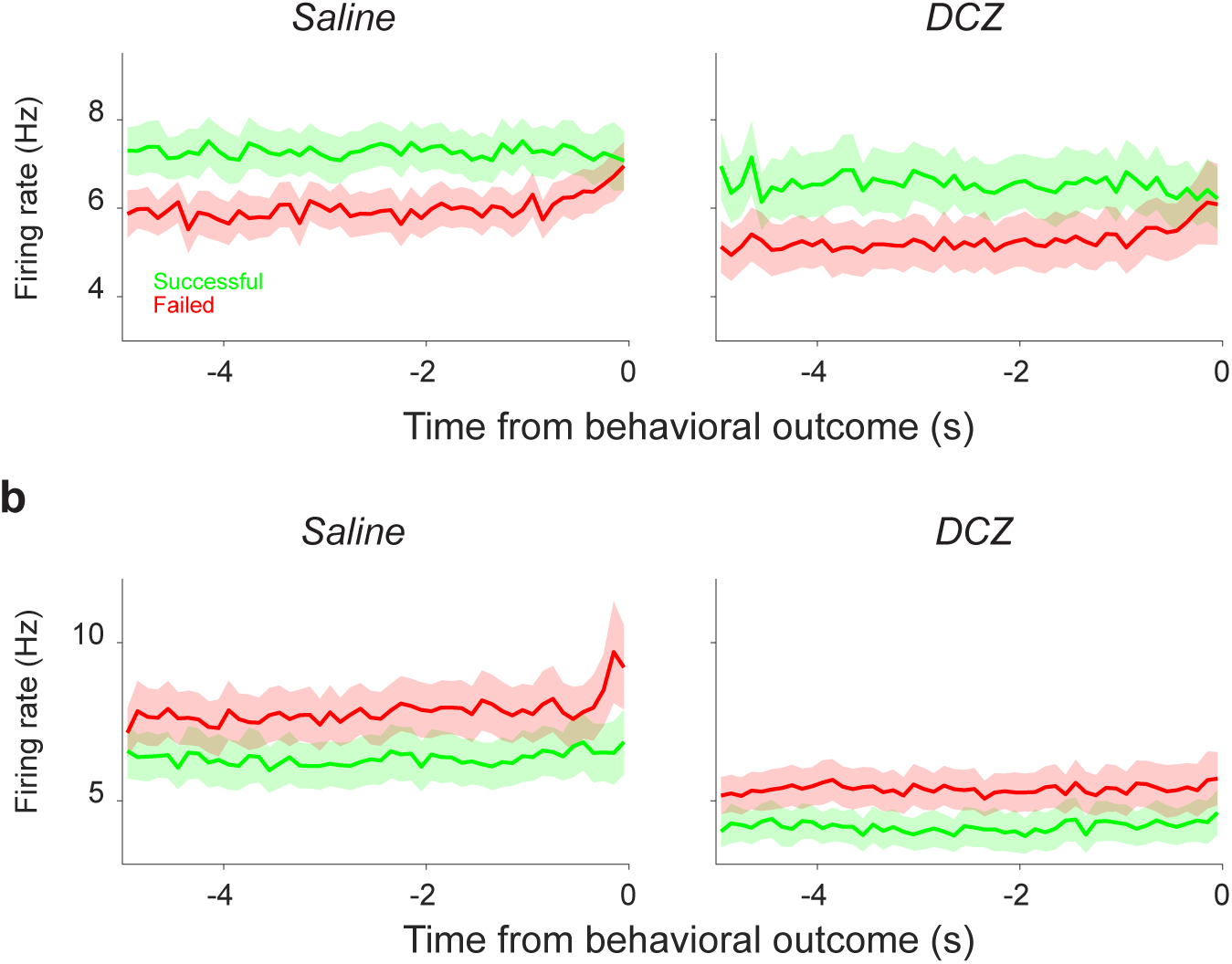
Firing rates of prefrontal encoding neurons during inhibitory control. **a)** Population firing rates prior to behavioral outcomes in the successful and failed trials under saline and DCZ treatment for encoding neurons with a higher firing rate in successful trials. **b)** Population firing rate prior to behavioral outcomes in the successful and failed trials under saline and DCZ treatment for encoding neurons with a lower firing rate in successful trials. All data are from 15 saline sessions and 15 DCZ sessions from 3 animals.

**Supplemental figure 11.**
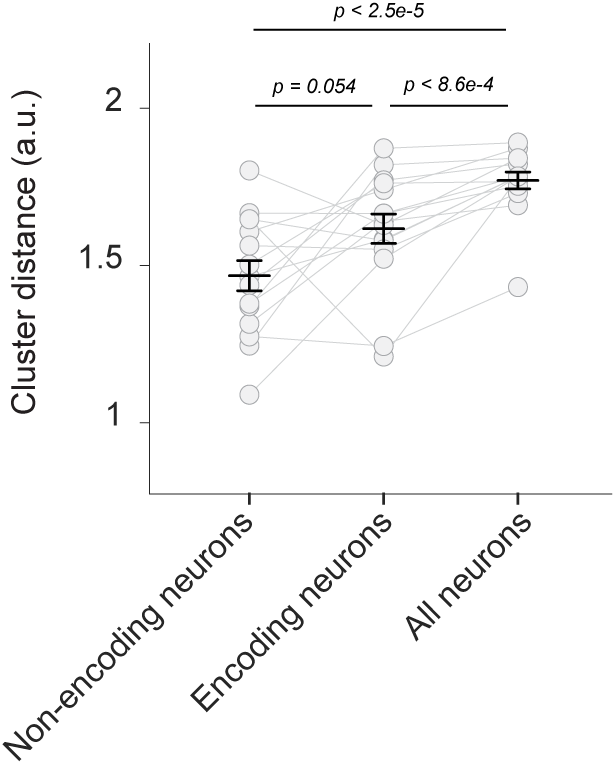
Non-encoding neurons in the prefrontal cortex contribute to population firing patterns representing inhibitory control.

**Supplemental figure 12.**
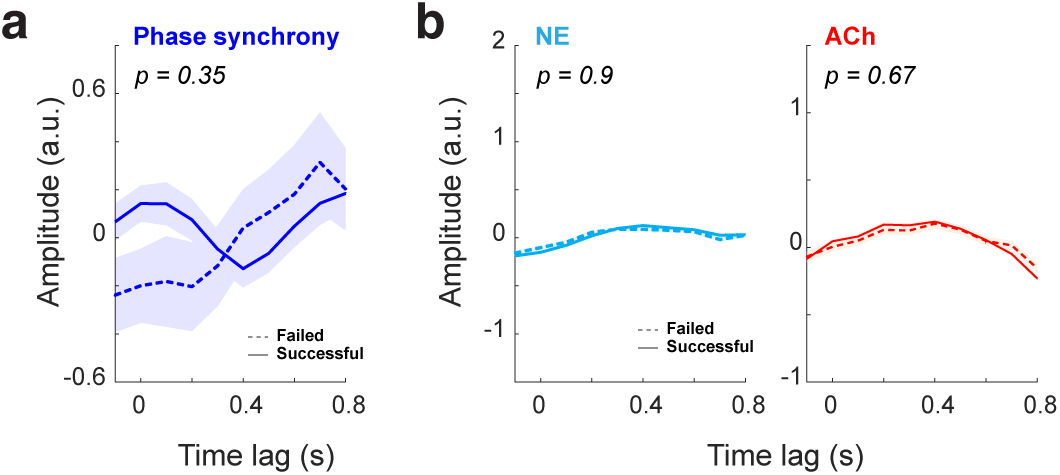
Temporal response functions (TRFs) during the free period. **a)** TRF mapping prefrontal NE-ACh phase synchrony to pupil size in successful and failed trials during the free period. **b)** TRF mapping NE (left) and ACh (right) signals to pupil size in successful and failed trials during the free period.

